# Chelator sensing and lipopeptide interplay mediates molecular interspecies interactions between soil bacilli and pseudomonads

**DOI:** 10.1101/2021.02.22.432387

**Authors:** Sofija Andric, Thibault Meyer, Augustin Rigolet, Anthony Argüelles Arias, Sébastien Steels, Grégory Hoff, Monica Höfte, René De Mot, Andrea McCann, Edwin De Pauw, Marc Ongena

## Abstract

Some bacterial species are important members of the rhizosphere microbiome and confer protection to the host plant against pathogens. However, our knowledge of the multitrophic interactions determining the ecological fitness of these biocontrol bacteria in their highly competitive natural niche is still limited. In this work, we investigated the molecular mechanisms underlying interactions between *B. velezensis,* considered as model plant-associated and beneficial species in the *Bacillus* genus, and *Pseudomonas* as a rhizosphere-dwelling competitor. Our data show that *B. velezensis* boosts its arsenal of specialized antibacterials upon the perception of the secondary siderophore enantio-pyochelin produced by phylogenetically distinct pseudomonads and some other genera. We postulate that *B. velezensis* has developed some chelator sensing systems to learn about the identity of its surrounding competitors. Illustrating the multifaceted molecular response of *Bacillus*, surfactin is another crucial component of the secondary metabolome mobilized in interbacteria competition. Its accumulation not only enhances motility but, unexpectedly, the lipopeptide also acts as a chemical trap that reduces the toxicity of other lipopeptides released by *Pseudomonas* challengers. This in turn favors the persistence of *Bacillus* populations upon competitive root colonization. Our work thus highlights new ecological roles for bacterial secondary metabolites acting as key drivers of social interactions.

---

Soil is one of the richest ecosystems in terms of microbial diversity and abundance^1^. However, the scarcity of resources makes it one of the most privileged environments for competitive interspecies interactions^2, 3^. A subset of the bulk soil microbes has evolved to dwell in the rhizosphere compartment surrounding roots due to continued nutrient-enriched exudation from the plant. Compared to bulk soil, microbial warfare in the rhizosphere is presumably even more intense as the habitat is spatially restricted and more densely populated^3^. Besides rivalry for nutrients (exploitative), interference competition is considered a key factor driving microbial interactions and community assembly. This competition can involve signal interference or toxins deployed by contact-dependent delivery systems^4, 5^ but is mainly mediated at distance through the emission of various molecular weapons. The molecular basis of interference interactions and their phenotypic outcomes between diverse soil bacterial species have been amply investigated in the last decade^6, 7^.

Bacilli belonging to the *B. subtilis* complex are ubiquitous members of the rhizosphere microbiome^2, 8, 9^. Among these species, *B. velezensis* has emerged as plant-associated model bacilli, displaying strong potential as biocontrol agent reducing diseases caused by phytopathogens^10^. *B. velezensis* distinguishes itself from other species of the *B. subtilis* group by its richness in biosynthetic gene clusters (BGCs, representing up to 13% of the whole genome) responsible for the synthesis of bioactive secondary metabolites (BSMs)^11, 12^. This chemically-diverse secondary metabolome includes volatiles, terpenes, non-ribosomal (NR) dipeptides, cyclic lipopeptides (CLPs) and polyketides (PKs), but also ribosomally synthesized lantibiotics and larger bacteriocins (RiPP)^13, 14^. BSMs are involved in biocontrol activity via direct inhibition of pathogenic microbes and/or via stimulation of the plant immune system^15, 16^. From an ecological viewpoint, BSMs also contribute to competitiveness in the rhizosphere niche thanks to multiple and complementary functions as drivers of developmental traits, as antimicrobials, or as signals initiating cross-talk with the host plant^17–19^.

Mostly guided by practical concerns for use as biocontrol agents, research on BSMs has mainly focused on the characterization of their biological activities. However, the impact of environmental factors that may modulate their expression under natural conditions still remains poorly understood. It includes interactions with other organisms sharing the niche. Some recent reports illustrate how soil bacilli may adapt their behavior upon sensing bacterial competitors but almost exclusively focusing on developmental traits (sporulation, biofilm formation, or motility)^7^.

Unlike other genera such as *Streptomyces*, it remains largely unknown to what extent bacilli in general and *B. velezensis* in particular, may modulate the expression of their secondary metabolome upon interaction with other bacteria^7, 20^. In this work, we investigated the molecular outcomes of interspecies interactions in which *B. velezensis* may engage. We selected *Pseudomonas* as challenger considering that species of this genus are also highly competitive and commonly encountered in rhizosphere microbiomes^8^. We performed experiments under nutritional conditions mimicking the oligotrophic rhizosphere environment and used contact-independent settings for pairwise interaction which probably best reflect the real situation in soil. Our data revealed that the two bacteria initiate multifaceted interactions mostly mediated by the non-ribosomally synthesized components of their secondary metabolome. We pointed out unsuspected roles for some of these BSMs in the interaction context. Beyond its role as a metal chelator, the *Pseudomonas* secondary siderophore enantio-pyochelin (E-PCH) acts as a signal triggering dual production of PKs and RiPP in *Bacillus,* while specific lipopeptides modulate the inhibitory interaction between the two species. This results in marked phenotypic changes in *B. velezensis* such as higher antibacterial potential, enhanced motility and protective effect via chemical trapping. We also illustrate the relevance of these outcomes in the context of competitive root colonization.

## Results

### *B. velezensis* modulates its secondary metabolome and boosts antibacterial activity upon sensing *Pseudomonas* metabolites

We used *B. velezensis* strain GA1 as a BSM-rich and genetically amenable isolate representative of the species. Genome mining with AntiSMASH 5.0^21^ confirmed the presence of all gene clusters necessary for the biosynthesis of known BSMs typically formed by this bacterium (Supplementary Table 1). Based on the exact mass and absence of the corresponding peaks in deletion mutants, most of the predicted non-ribosomal (NR) secondary metabolites were identified in cell-free crude supernatants via optimized UPLC-MS (Supplementary Fig. 1). It includes the whole set of cyclic lipopeptides (CLPs of the surfactin, fengycin and iturin families) and polyketides (PKs difficidin, macrolactin and bacillaene) with their multiple co-produced structural variants, as well as the siderophore bacillibactin. We verified that all these compounds are readily formed in the so-called exudate-mimicking (EM) medium reflecting the specific content in major carbon sources typically released by roots of *Solanaceae* (such as tomato) plants^22^. In addition to these NR products, genes encoding RiPPs such as amylocyclicin and amylolysin are also present in GA1, but these compounds could not be reliably detected in culture broths. The NR dipeptide bacilysin was also predicted but not detected. We selected as the main interaction partner the plant-associated *Pseudomonas* sp. strain CMR12a based on its biocontrol potential and its production of multiple secondary metabolites^23–27^. Genome mining confirmed the potential of CMR12a to synthesize a range of BSMs, including antimicrobial phenazines, the siderophores pyoverdine (PVD) (structure confirmation in Supplementary Fig. 2) and E-PCH as well as two structurally distinct CLPs, sessilins and orfamides (Supplementary Table 1). In contrast to *Bacillus*, the capacity to co-produce two different CLPs is a quite rare trait for non-phytopathogenic pseudomonads and represented an additional criterion for selecting strain CMR12a for this study^23, 27–29^. In the case of CMR12a, according to the exact mass and absence of the corresponding peaks in deletion mutants, all these compounds were detected in EM culture broth but most of them are more efficiently produced upon growth in casamino acids medium (CAA) commonly used for *Pseudomonas* cultivation (Supplementary Fig. 1).

Our prime objective was to evaluate the intrinsic potential of *B. velezensis* to react to the perception of *Pseudomonas* metabolites in an experimental setting avoiding interferences due to diffusion constraints in a semi-solid matrix or due to the formation of impermeable biofilm structures. The first assays were performed by growing GA1 in agitated liquid EM medium supplemented or not with (sterile) BSM-containing spent medium of CAA-grown CMR12a (CFS, cell-free supernatant). At a low dose (2% (v/v)), the addition of this CFS extract led to a marked increase in the production of some GA1 NR metabolites. Significantly higher amounts were measured for surfactins, bacillaene or its dehydrated variant dihydrobacillaene (2H-bae), difficidin or its oxidized form, and bacillibactin (Fig. 1a, b) but not for other compounds such as fengycins, iturins and macrolactins (Fig. 1a).

**Figure 1.**
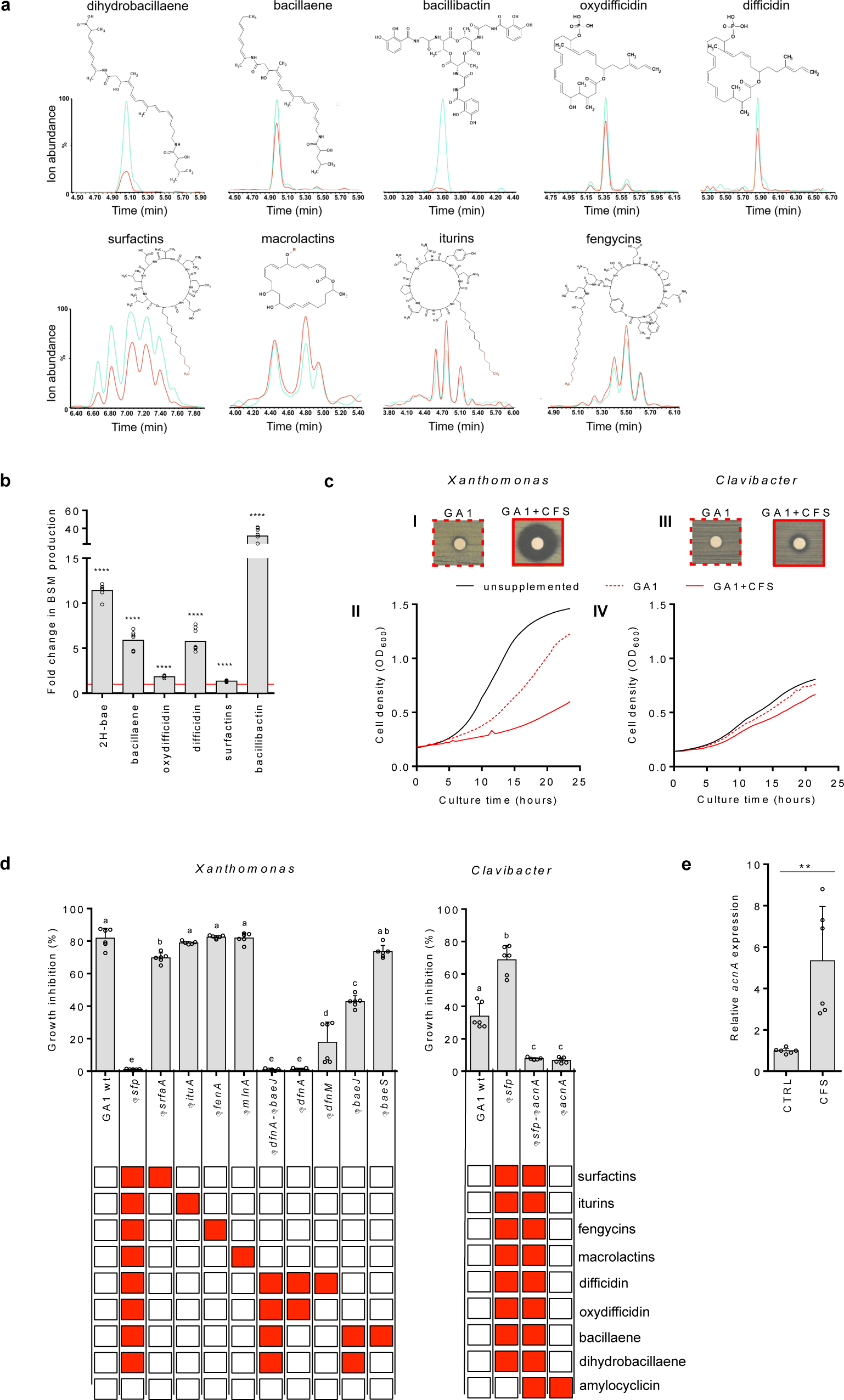
Stimulation of BSMs production by *B. velezensis* GA1 and enhanced anti-bacterial activities in response to *Pseudomonas* sp. CMR12a secreted metabolites. **a.** UPLC-MS extracted ions chromatograms (EIC) illustrating the relative abundance of ions corresponding to non-ribosomal metabolites produced by *B. velezensis* GA1 in CFS-supplemented (2% v/v) EM medium (blue) compared to un-supplemented cultures used as control (red). Red-coloured parts in the representation of lipopeptides and macrolactin illustrate the variable structural traits explaining the occurrence of naturally co-produced variants (multiple peaks) **b**. Fold increase in GA1 BSM production upon addition of CMR12a CFS (2% v/v) compared to un-supplemented cultures (fold change = 1, red line). Data were calculated based on the relative quantification of the compounds by UPLC-MS (peak area) in both conditions. Mean values were calculated from data obtained in three cultures (repeats) from two independent experiments (n = 6). Statistical significance was calculated using Mann–Whitney test where ‘’****’’ represents significant difference at *P*<0.0001. **c.** Enhanced Anti-*Xanthomonas campestris* (**I** and **II**) and anti-*Clavibacter michiganensis* (**III** and **IV**) activities of GA1 extracts (cell-free culture supernatant) after growth in CMR12a CFS-supplemented medium (GA1+CFS) compared to control (GA1). It was assessed both on plates by the increase in inhibition zone around paper disc soaked with 5 µl the GA1 extracts (**I** and **III**) and in liquid cultures of the pathogens by reduction of growth upon addition of 4% (v/v) of GA1 extracts (**II** and **IV**). Data are from one representative experiment and similar results were obtained in two independent replicates. **d.** Antibacterial activities of extracts from GA1 WT and mutants impaired in production of specific BSMs. Metabolites not produced by the different mutants are illustrated with red boxes in the table below. All values represent means with error bars indicating SD calculated on data from three cultures (repeats) in two independent experiments (n = 6). Letters a to d indicate statistically significant differences according to one-way analysis of variance (ANOVA) and Tukey’s HSD test (Honestly significantly different, α = 0.05). **e.** Differential expression of the *acnA* gene encoding the amylocylicin precursor, upon supplementation with CMR12a CFS compared to GA1 un-supplemented culture. Mean and SD values, n = 6, “**” indicates statistical significance according to Mann–Whitney test, *P*<0.01.

The boost in BSMs synthesis triggered by *Pseudomonas* CFS was associated with an increase in the antibacterial activity of the corresponding extracts when tested for growth inhibition of *Xanthomonas campestris* and *Clavibacter michiganensis* used respectively as representative of Gram-negative and Gram-positive plant pathogenic bacteria of agronomical importance^30^ (Fig. 1c). Since most of the BSMs are not commercially available and our attempt to purify PKs failed due to chemical instability, we could not use individual compounds for their specific involvement in bacterial inhibition. As an alternative, we generated and tested a range of GA1 knockout mutants including the Δ*sfp* derivative specifically repressed in 4’-phosphopantetheinyl transferase essential for the proper functioning of the PK and NRP biosynthesis machinery. Full loss of anti- *Xanthomonas* activity in Δ*sfp* extracts indicated a key role for NR BSMs and ruled out the possible involvement of other chemicals known for their antibacterial activity such as bacilysin or RiPPs (Fig. 1d). Loss of function of mutants specifically repressed in the synthesis of individual compounds pointed out the key role of (oxy)difficidin and to a lower extent of 2H-bae in *Xanthomonas* inhibition (Fig. 1d). These two PKs are also responsible for GA1 inhibitory activity toward other important bacterial phytopathogens such as *Pectobacterium carotovorum*, *Agrobacterium tumefaciens* and *Rhodococcus fasciens* but are not involved in the inhibition of plant pathogenic *Pseudomonas* species for which bacilysin may be the active metabolite (Supplementary Fig. 3). However, as illustrated below, *B. velezensis* does not display significant toxicity against CMR12a and other non-pathogenic soil *Pseudomonas* isolates tested here. Stimulation of PKs synthesis upon sensing CMR12a is not specific to GA1 and was also observed in other *B. velezensis* strains with well-known biocontrol potential such as S499, FZB42 and QST713^31–33^ (Supplementary Fig. 4).

In contrast to *Xanthomonas*, enhanced antibiotic activity against *Clavibacter* is not mediated by NR products as shown by the fully conserved activity in the Δ*sfp* mutant (Fig. 1d). Therefore, we suspected from genomic data and literature^34^ that RiPPs such as amylocyclicin could be involved in inhibition. This hypothesis was supported by the 80% reduction in antibiotic potential observed for the Δ*acnA* mutant knocked out for the corresponding biosynthesis gene (Fig. 1d). Besides, RT-qPCR data revealed a highly induced expression of *acnA* gene in GA1 cells upon supplementation with CMR12a CFS (Fig. 1e). However, we were not able to provide evidence for higher accumulation of the mature peptide in the medium. Enhanced expression of the *acnA* gene in presence of *Pseudomonas* products was also observed for strain S499 (Supplementary Fig. 5).

### E-PCH acts as a signal sensed by *Bacillus* to stimulate polyketide production

We next wanted to identify the signaling molecules secreted by *Pseudomonas* that are sensed by *Bacillus* cells and lead to improved BSMs production. For that purpose, we used 2H-bae as an indicator of the *Bacillus* response because it represents the most consistent and highly boosted polyketide. We first compared the triggering potential of CFS obtained from knockout mutants of CMR12a specifically lacking the different identified metabolites (Supplementary Fig. 1). Only extracts from mutants impaired in the production of siderophores and more specifically E-PCH were significantly affected in PKs-inducing potential (Fig. 2a). Possible involvement of this compound was supported by the drastic reduction in the activity of CFS prepared from CMR12a culture in CAA medium supplemented with Fe^3+^ where siderophore expression is repressed (Fig. 2b, Supplementary Fig. 6). We also performed bioactivity-guided fractionation and data showed that only extracts containing PVD and/or E-PCH displayed consistent PKs-triggering activity (Supplementary Fig. 7). HPLC-purified compounds were also tested independently at a concentration similar to the one measured in CFS CAA extract revealing a much higher PK-triggering activity for E-PCH compared to the main PVD isoform (Fig. 2b). Dose-dependent assays further indicated that supplementation with PVD, as strong chelator^35^, caused iron limitation in the medium which is sensed by GA1. It is supported by the marked increase in production of the siderophore bacillibactin in GA1 wild-type (Fig. 2c) and by the reduced growth of the Δ*dhbC* mutant, repressed in bacillibactin synthesis, upon PVD addition (Fig. 2d, Supplementary Fig. 8). This last result indicates that PVD in its ferric form cannot be taken up by GA1 despite the presence of several transporters for exogenous siderophores in *B. velezensis* similar to those identified in *B. subtilis*^36, 37^ based on genome comparison (Supplementary Table 2). Therefore, we assumed that iron stress mediated by PVD only induces a rather limited boost in PKs production. We validated that such response is not due to iron starvation by supplementing GA1 culture with increasing doses of the 2,2’-dipyridyl (DIP) chemical chelator that cannot be taken up by *Bacillus* cells (Fig. 2b). By contrast, the addition of E-PCH with a much lower affinity for iron does not activate bacillibactin synthesis (Fig. 2c) and does not affect Δ*dhbC* growth at the concentrations used (Fig. 2d, Supplementary Fig. 8). We conclude that the activity of this compound referred to as secondary siderophore is not related to iron-stress. If internalized, E-PCH can cause oxidative stress and damage in other bacteria as reported for *E. coli*^38, 39^. However, the absence of toxicity toward GA1 indicates that E-PCH is not taken up by *Bacillus* cells and thus clearly acts as a signal molecule perceived at the cell surface. PKs boost also occurred upon addition of CFS obtained from other *Pseudomonas* isolates producing pyochelin-type siderophores, such as *P. protegens* Pf-5^40^. However, PKs stimulation was similarly observed in response to *P. tolaasii* CH36^41^ which does not form pyochelin, indicating that other unidentified BSMs may act as triggers in other strains (Supplementary Fig. 9).

**Figure 2:**
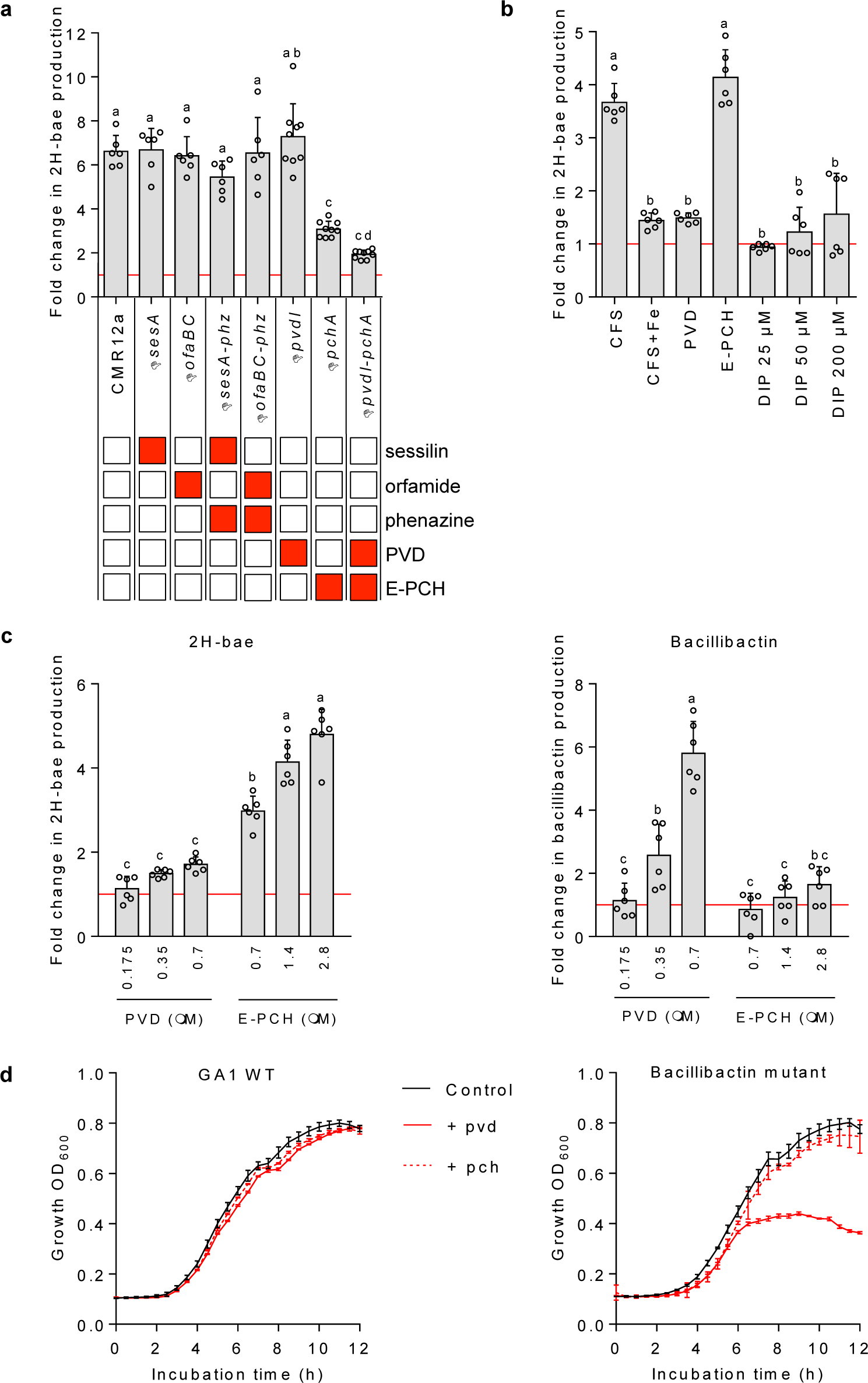
E-PCH as main *Pseudomonas* trigger of anti-bacterial activity boosted in *B. velezensis* GA1. **a**, Effect of GA1 culture supplementation with CFS (2% v/v) from CMR12a WT and various mutants on dihydrobacillaene (2H-bae) production. Metabolites specifically repressed in the different CMR12a mutants are illustrated by red boxes. Fold changes were calculated based on relative quantification of the compounds by UPLC-MS (peak area) in treated cultures compared to un-supplemented controls (fold change = 1, red line). Data are means and SE calculated from three replicate cultures in two (n = 6) or three (n = 9) independent experiments and different letters indicate statistically significant differences (ANOVA and Tukey’s test, α = 0.05). **b,** Differential production of 2H-bae after addition of 0.35 µM pure PVD, 1.4 µM pure E-PCH, 4% v/v *Pseudomonas* sp. CMR12a CFS (CFS CAA), CMR12a CFS from iron supplemented culture (CFS CAA+Fe) and different concentration of the iron-chelating agent 2,2’-dipyridyl (DIP). Data are expressed and were statistically treated as described in **a** with n = 6 in all treatments. **c,** Dose-dependent effect of pure PVD and E-PCH on bacillibactin and 2H-bae production. GA1 cultures were supplemented with the indicated concentrations of HPLC-purified CMR12a siderophores. Experiments were replicated and data statistically processed as described in **b**. **d,** Impact of the addition of pure PVD and E-PCH on the growth of GA1 WT and its Δ*dhbC* mutant repressed in bacillibactin synthesis. *Pseudomonas* siderophores were added at a final concentration similar to the one obtained by adding CMR12a CFS at 4% v/v. Means and SD are from three replicates. See Supplementary Figure 8 for detailed data and statistical significance.

### Enhanced motility as distance-dependent and surfactin-mediated response of *Bacillus*

Surfactin production is stimulated by CMR12a CFS and by pure E-PCH (Fig. 1a and Supplementary Fig. 10). Based on mutant loss-of-function analysis, this multifunctional CLP does not contribute to the antibacterial potential of *B. velezensis* (Fig. 1d and Supplementary Fig. 3) but is known to be notably involved in developmental processes of multicellular communities such as biofilm formation and motility^42^. Therefore, we wanted to test a possible impact of *Pseudomonas* on the motile phenotype of *B. velezensis* upon co-cultivation on plates. We observed distance-dependent enhanced motility on medium containing high agar concentrations (1.5% m/v) which phenotypically resembles the sliding-type of motility illustrated by typical “van Gogh bundles”^42^ (Fig. 3a). This migration pattern is flagellum independent but depends on multiple factors including the synthesis of surfactin which reduces friction at the cell-substrate interface^42^. We thus suspected such enhanced motility to derive from an increased formation of the lipopeptide. This was supported by the almost full loss of migration of the Δ*srfaA* mutant in these interaction conditions (Fig. 3b). Moreover, spatial mapping via MALDI-FT-ICR MS imaging confirmed a higher accumulation of surfactin ions in the interaction zone and around the *Bacillus* colony when growing at a short or intermediate distance from the *Pseudomonas* challenger, compared to the largest distance where the motile phenotype is much less visible (Fig. 3c). These data indicate that *Bacillus* cells in the microcolony perceive a soluble signal diffusing from the *Pseudomonas* colony over a limited distance.

**Figure 3:**
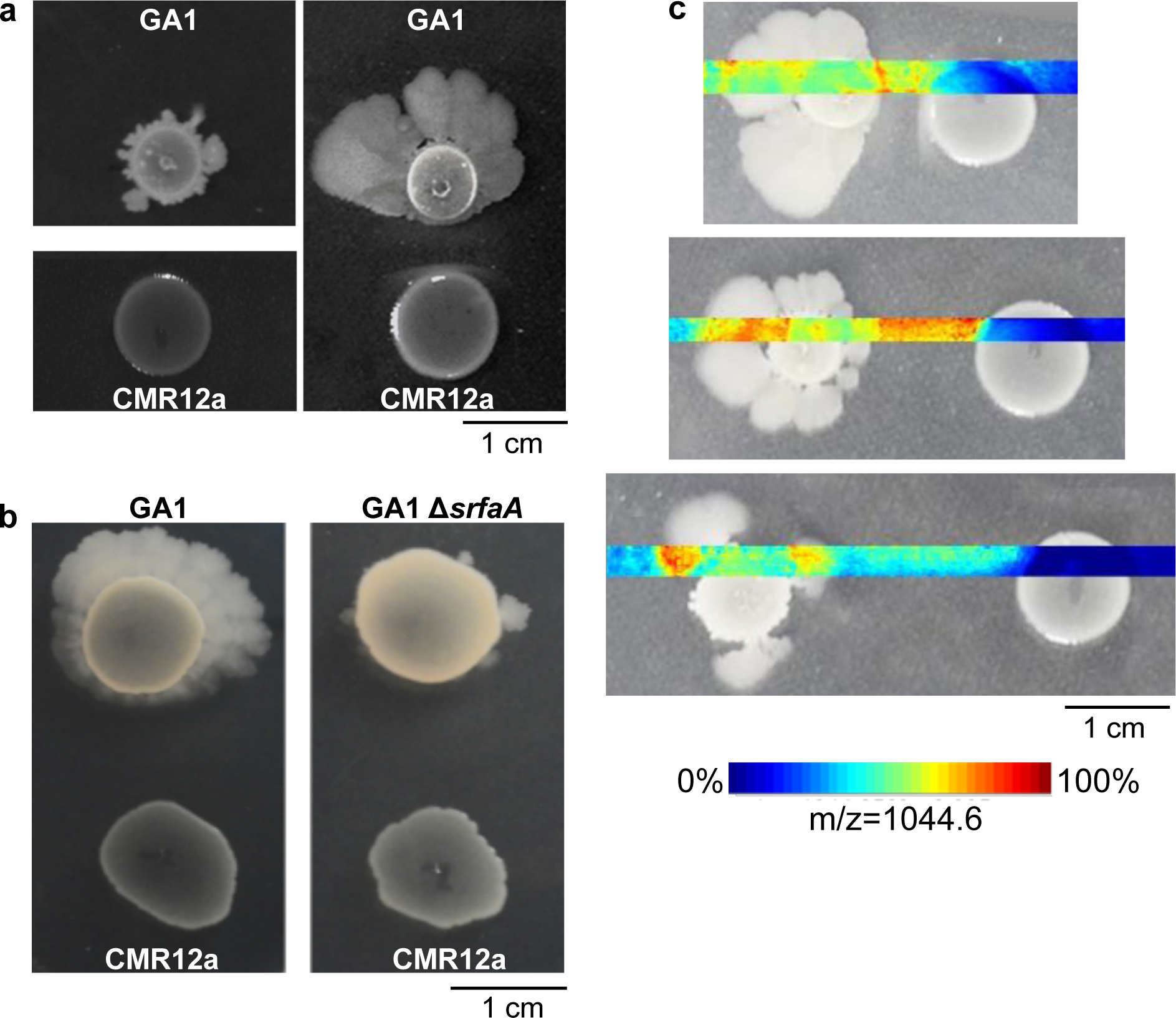
Distance- and surfactin-dependent enhanced motility of *B. velezensis* GA1 in interaction with *Pseudomonas* CMR12a. **a,** GA1 motility phenotype on EM gelified medium when cultured alone (left panel) or in confrontation with CMR12a at a short distance (1 cm) (right panel). **b,** Motility pattern of GA1 or his Δ*srfaA* surfactin deficient mutant in confrontation with CMR12a at a short distance. **c,** MALDI FT-ICR MSI (Mass spectrometry Imaging) heatmaps showing spatial localization and relative abundance of ions ([M+Na]^+^) corresponding to the C_14_ surfactin homolog (most abundant) when *B. velezensis* GA1 is in confrontation with CMR12a at increasing distances (one biological replicate).

### Interplay between CLPs drives antagonistic interactions

Besides modulating secondary metabolite synthesis, we further observed that confrontation with *Pseudomonas* may also lead to some antagonistic outcomes. GA1 growth as planktonic cells is slightly inhibited upon supplementation of the medium with 2% v/v CMR12a CFS but this inhibition is much more marked at a higher dose (Supplementary Fig. 11). To identify the *Pseudomonas* compound retaining such antibiotic activity, we tested the effect of CFS from various CMR12a mutants impaired in the synthesis of lipopeptides and/or phenazines. Even if some contribution of other compounds cannot be ruled out, it revealed that the CLP sessilin is mainly responsible for toxic activity toward GA1 grown in liquid cultures but also when the two bacteria are grown at close proximity on gelified EM medium (Fig. 4a). Nevertheless, we observed that the sessilin-mediated inhibitory effect is markedly reduced by delaying CFS supplementation until 6 h of *Bacillus* culture instead of adding it at the beginning of incubation (Fig. 4b). This suggested that early secreted *Bacillus* compounds may counteract the toxic effect of sessilin. We hypothesized that surfactin can play this role as it is the first detectable BSM to accumulate in significant amounts in the medium early in the growth phase. We tested the surfactin-deficient mutant in the same conditions and observed that its growth is still strongly affected indicating that no other GA1 compound may be involved in toxicity alleviation. Chemical complementation with purified surfactins restored growth to a large extent, providing further evidence for a protective role of the surfactin lipopeptide (Fig. 4b). Such sessilin-dependent inhibition also occurred when bacteria were confronted on solid CAA medium (Fig. 4c-I) favoring *Pseudomonas* BSM production. In these conditions, the formation of a white precipitate in the interaction zone was observed with CMR12a wild-type but not when GA1 was confronted with the Δ*sesA* mutant (Fig. 4c). UPLC-MS analysis of ethanol extracts from this white-line area confirmed the presence of sessilin ions but also revealed an accumulation of surfactin from GA1 in the confrontation zone (Fig. 4d). The involvement of surfactins in precipitate formation was confirmed by the absence of this white-line upon testing the Δ*srfaA* mutant of GA1 (Fig. 4c-I, II). The loss of surfactin production and white-line formation was associated with a higher sensitivity of the *Bacillus* colony to the sessilin toxin secreted by *Pseudomonas*. Altogether, these data indicate that surfactin acts as a chemical trap and inactivates sessilin via co-aggregation into insoluble complexes.

**Figure 4:**
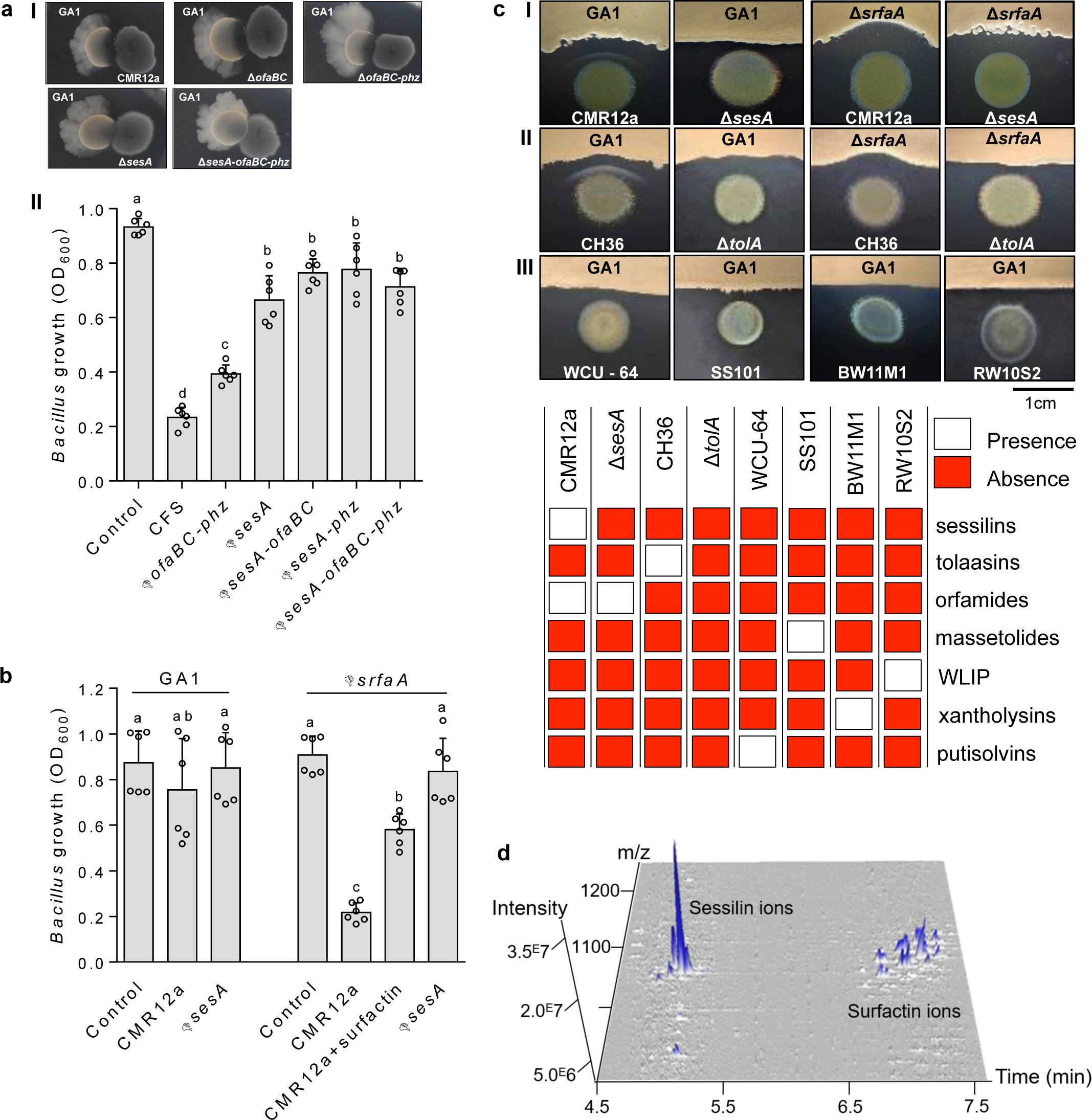
Surfactin attenuates sessilin-mediated toxicity via white-line formation. **a,** I. Polarized inhibition of GA1 micro-colony development upon co-cultivation at close contact with CMR12a colonies on EM plates. **II.** Inhibition of GA1 cell growth in EM liquid culture supplemented with 6% v/v of CFS prepared from CMR12a wild-type or mutants repressed in the synthesis of orfamides and phenazines (Δ*ofaBC-phz*), sessilins (Δ*sesA*), sessilins and orfamides (Δ*sesA-ofaBC*), sessilins and phenazines (Δ*sesA-phz*), or of all compounds (Δ*sesA-ofaBC-phz*). Data show mean and SD calculated from two independent experiments each with three culture replicates (n = 6) and different letters indicate statistically significant differences (ANOVA and Tukey’s test, α = 0.05). **b,** Growth inhibition of GA1 WT and Δ*srfaA* mutant upon delayed supplementation (added 6 h after incubation start) with CFS from CMR12a WT alone or together with pure surfactin as chemical complementation) and with CFS from the sessilin mutant (Δ*sesA*). Un-supplemented cultures of GA1 were used as control. Experiments were replicated and data statistically processed as described in **a**. **c,** White line formation and/or *Bacillus* inhibition observed upon confrontation of GA1 WT or the surfactin mutant Δ*srfaA* with (**I**) CMR12a or its Δ*sesA* derivative, (**II**) *P. tolaasii* CH36 or its tolaasin defective mutant Δ*tolA* and (**III**) other *Pseudomonas* CLP producers WCU-84, SS101, BW11M1, RW10S2. CLPs produced by the individual *Pseudomonas* strains are mentioned in the chart below. **d,** 3D representation of UPLC-MS analysis of CLPs that are present in the white-line zone between GA1 and CMR12a (**I**). It shows the specific accumulation of sessilin and surfactin molecular ions (one biological replicate).

A similar CLP-dependent antagonistic interaction and white-line formation were observed upon co-cultivation of GA1 with *P. tolaasii* strain CH36 producing tolaasin (Fig. 4c-II), a CLP structurally very similar to sessilin (only differing by two amino acid residues, Supplementary Fig. 12). However, this chemical aggregation is quite specific regarding the type of CLP involved, since it was not visible upon the interaction of GA1 with other *Pseudomonas* strains forming different CLP structural groups that are not toxic for *Bacillus* (Fig. 4c-III, see Supplementary Fig. 12 and 13 for identification and structures). Sessilin/tolaasin-dependent toxicity and white-line formation were also observed when other surfactin-producing *B. velezensis* isolates were confronted with CMR12a and CH36 (Supplementary Fig. 14 and 15, respectively). Although the chemical basis and the stoichiometry of such molecular interaction remain to be determined, it probably follows the same rules as observed for the association between sessilins/tolaasins and other endogenous *Pseudomonas* CLPs such as WLIP or orfamides^28^ or between CLPs and other unknown metabolites^43, 44^.

### BSMs-mediated interactions drive competitive root colonization

Our *in vitro* data point out how *B. velezensis* may modulate its secondary metabolome when confronted with *Pseudomonas*. To appreciate the relevance of our findings in a more realistic context, we next evaluated whether such BSMs interplay may also occur upon root co-colonization of tomato plantlets and possibly impact *Bacillus* fitness. When inoculated independently, CMR12a colonized roots more efficiently than GA1 within the first 3 days, most probably due to a higher intrinsic growth rate^45^. Upon co-inoculation, the CMR12a colonization rate was not affected but GA1 populations were reduced compared to mono-inoculated plantlets (Fig. 5a). UPLC-MS analysis of methanolic extracts prepared from co-bacterized roots (and surrounding medium) revealed substantial amounts of E-PCH (Supplementary Fig. 16) indicating that the molecule is readily formed under these conditions and could therefore also act as a signal *in planta.* Probably due to the low populations of GA1, we could not detect *Bacillus* PKs and RiPPs in these extracts. However, a significantly enhanced expression of gene clusters responsible for the synthesis of bacillaene, difficidin and amylocyclicin was observed in GA1 cells co-inoculated with *Pseudomonas* compared to single inoculation (Fig. 5b). It indicated that the metabolite response observed in GA1 *in vitro* cultures in EM medium may also occur upon competitive colonization where the bacteria feed exclusively on root exudates.

**Figure 5:**
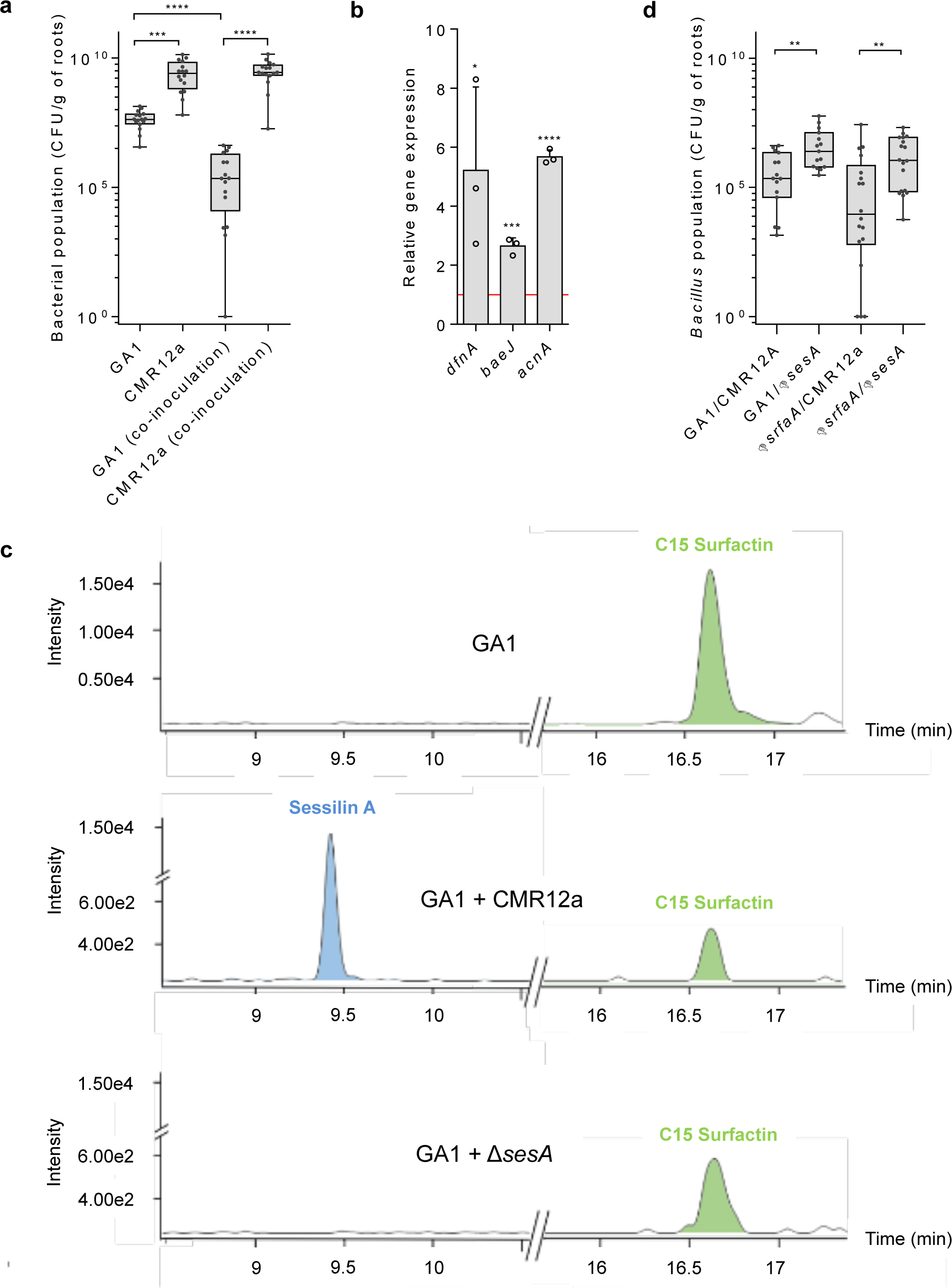
Competitive colonization assays support the roles of BSMs in *Bacillus-Pseudomonas* interaction *in planta*. **a,** GA1 and CMR12a cell populations as recovered from roots at 3 days post-inoculation (dpi) of tomato plantlets when inoculated alone (GA1, CMR12a) or co-inoculated (co-inoculation). Box plots were generated based on data from three independent assays each involving at least 4 plants per treatment (n=16). The whiskers extend to the minimum and maximum values, and the midline indicates the median. Statistical differences between the treatments were calculated using Mann–Whitney test and ‘’****’’ and ‘’***’’ represent significant differences at *P*<0.0001 and *P*<0.001, respectively. **b,** *In planta* (3 dpi on tomato roots) relative expression of the *dfnA, baeJ* and *acnA* genes responsible for the synthesis of respectively (oxy)difficidin, 2H-bae and amylocyclicin. Graphs show the mean and SD calculated from three biological replicates (n = 3) each involving six plants. Fold change = 1 as red line corresponds to gene expressions in GA1 inoculated alone on roots used as control conditions. Statistical comparison between data in co-colonization setting and control conditions was performed based on T-test (*, *P*<0.05; **, *P*<0.01; ***, *P*<0.001; ****, *P*<0.0001). **c.** UPLC-MS EIC illustrating relative *in planta* production of sessilins and surfactins by monocultures of *B. velezensis* GA1 (GA1) and co-cultures of wild-types (GA1+CMR12a) and *B. velezensis* GA1 and *Pseudomonas* sp. CMR12a impaired in sessilins production (GA1+Δ*sesA*). **d,** Cell populations recovered at 3 dpi for GA1 WT (GA1) or the surfactins impaired mutant (Δ*srfaA*) co-inoculated with CMR12a WT (CMR12a) or its sessilins KO mutant (Δ*sesA*). See **a** for replicates and statistics (**, *P*<0.01).

Lipopeptides involved in interference interaction are also readily formed upon single and dual root colonization (Fig. 5d). We hypothesized that the inhibitory effect of sessilin may impact the colonization potential of GA1 in presence of CMR12a which was confirmed by the increase in GA1 populations co-inoculated with the Δ*sesA* mutant (Fig. 5c). Moreover, colonization by the Δ*srfaA* mutant is more impacted compared to WT when co-cultivated with CMR12a and a significant gain in root establishment is recovered upon co-colonization with the Δ*sesA* mutant (Fig. 5d). The sessilin-surfactin interplay thus also occurs *in planta.* Sessilin would confer a competitive advantage to CMR12a during colonization by inhibiting GA1 development but efficient surfactin production on roots may provide some protection to the *Bacillus* cells.

## Discussion

It has been recently reported that *Pseudomonas* toxin delivery via Type VI secretion system and antibiotic (2,4-diacetylphloroglucinol) production may impact biofilm formation and sporulation in *B. subtilis*^46, 47^. However, our current understanding of the molecular basis of interactions between soil bacilli and pseudomonads is still rather limited. Here we show that the model species *B. velezensis* can mobilize a substantial part of its secondary metabolome in response to *Pseudomonas* competitors. To our knowledge, it is the first evidence for enhanced synthesis of both broad-spectrum polyketides and RiPP in *Bacillus* upon a perception of other bacteria, in contact-independent *in vitro* settings and upon competitive root colonization. This correlates with an enhanced antibacterial potential which is of interest for biocontrol but which can also be considered as a defensive strategy to persist in its natural competitive niche. Upon sensing *Pseudomonas*, *B. velezensis* calls on its antibiotic arsenal but also recruits its surfactin lipopeptide to improve multicellular mobility. This may be viewed as an escape mechanism enabling *Bacillus* cells to relocate after detecting harmful challengers. Improved motility of *B. subtilis* has been already described upon sensing competitors such as *Streptomyces venezuelae*^7, 48^ but no relationship was established with enhanced production of BSMs potentially involved in the process. We also highlight a new role for surfactin acting as a chemical shield to counteract the toxicity of exogenous CLPs. Intraspecies CLP co-precipitation has been reported^23^ but our results make sense of this phenomenon in the context of interference interaction between two different genera. *In planta*, this new function of surfactin contributes to *Bacillus* competitiveness for root invasion. This has to be added to other previously reported implications of surfactin in *B. subtilis* interspecies interactions, such as interfering with the growth of closely related species in synergy with cannibalism toxins^49^, inhibiting the development of *Streptomyces* aerial hyphae^50^, or participating in the expansion and motility of the interacting species^47^. We postulate that such *Bacillus* metabolite response largely contributes to mount a multi-faceted defensive strategy in order to gain fitness and persistence in its natural competitive niche.

Furthermore, we exemplify that PKs stimulation in *B. velezensis* is mainly mediated by the *Pseudomonas* secondary siderophore pyochelin, although it cannot be excluded that other secreted products may also play a role. *Bacillus* perceives pyochelin in a way independent of iron stress and piracy, indicating that beyond its iron-scavenging function, this siderophore may also act as infochemical in interspecies cross-talk. In the pairwise system used here, E-PCH signaling superimposes the possible effect of iron limitation in the external medium which may also result in enhanced production of antibacterial metabolites by *Bacillus,* as occasionally reported^51^. That said, due to the limitation in bioavailable iron, almost all known rhizobacterial species have adapted to produce their iron-scavenging molecules to compete for this essential element^52–54^. Siderophore production is thus widely conserved among soil-borne bacteria^55^. It means that upon recognition of exogenous siderophores, any isolate may somehow identify surrounding competitors. However, some of these siderophores are structurally very variable and almost strain-specific (such as PVDs from fluorescent pseudomonads) while some others are much more widely distributed across species and even genera (enterobactin-like, citrate)^52^. In both cases, their recognition would not provide proper information about the producer because they are too specific or too general, respectively. Interestingly, the synthesis of E-PCH and its structurally very close enantio form is conserved in several but not all *Pseudomonas* sp.^56–58^ as well as in a limited number of species belonging to other genera such as *Burkholderia*^59^ and *Streptomyces*^60, 61^. We therefore hypothesize that *Bacillus* may have evolved some chelator-sensing systems targeting siderophores that are conserved enough to be detected but restricted to specific microbial phylogenetic groups. With this mechanism, soil bacilli would rely on siderophores as public goods to accurately identify competitors and respond in an appropriate way like remodeling its BSM secretome. This novel concept of chelator sensing represents a new facet of siderophore-mediated social interactions. Whether it is used for other secondary siderophores than E-PCH and if so, whether this adaptative trait can be generalized to other soil-dwelling species deserves to be further investigated given its possible impact on soil bacterial ecology. Beyond the notion of specialized metabolites, we point out unsuspected functions for some bacterial small molecules in the context of interactions between clades that are important members of the plant-associated microbiome.

## Methods

### Bacterial strains and growth conditions

Strains and plasmids used in this study are listed in Supplementary Table 3. *B. velezensis* strains were grown at 30 °C on, half diluted, recomposed exudate solid medium (EM)^22^ or in liquid EM with shaking (160 rpm). Deletion mutants of *B. velezensis* were selected on appropriate antibiotics (chloramphenicol at 5 µg/ml, phleomycin at 4 µg/ml, kanamycin at 25 µg/ml) on Lysogeny broth (LB) (10 g l^-1^ NaCl, 5 g l^-1^ yeast extract and 10 g l^-1^ tryptone). *Pseudomonas* sp. strains were grown on King B (20 g l^-1^ of bacteriological peptone, 10 g l^-1^ of glycerol and 1.5 g l^-1^ of K_2_HPO_4_, 1.5 g l^-1^ of MgSO_4_.7H_2_O, pH = 7) and casamino acid (CAA) solid and liquid medium (10 g l^-1^ casamino acid, 0.3 g l^-1^ K2HPO4, 0.5 g l^-1^ MgSO4 and pH = 7) with shaking (120 rpm), at 30 °C. The phytopathogenic bacterial strains were grown on LB and EM solid and liquid media and with shaking (150 rpm), at 30 °C.

Construction of deletion mutants of *B. velezensis* GA1

All deletion mutants were constructed by marker replacement. Briefly, 1 kb of the upstream region of the targeted gene, an antibiotic marker (chloramphenicol, phleomycin or kanamycin cassette) and downstream region of the targeted gene were PCR amplified with specific primers (Supplementary Table 3). The three DNA fragments were linked by overlap PCR to obtain a DNA fragment containing the antibiotic marker flanked by the two homologous recombination regions. This latter fragment was introduced into *B. velezensis* GA1 by natural competence induced by nitrogen limitation^62^. Homologous recombination event was selected by chloramphenicol resistance (phleomycin resistance for double mutants or kanamycin resistance for triple mutants) on LB medium. All gene deletions were confirmed by PCR analysis with the corresponding UpF and DwR specific primers and by the loss of the corresponding BSMs production.

Transformation of the *B. velezensis* GA1 strain was performed following the protocol previously described^62^ with some modifications. One fresh GA1 colony was inoculated into LB liquid medium at 37 °C (160 rpm) until reaching an OD600_nm_ of 1.0. Afterwards, cells were washed one time with peptone water and one time with a modified Spizizen minimal salt liquid medium (MMG) (19 g l^-1^ K_2_HPO_4_ anhydrous; 6 g l^-1^ KH_2_PO_4_; 1 g l^-1^ Na_3_ citrate anhydrous; 0.2 g l^-1^ MgSO_4_ 7H_2_O; 2 g l^-1^ Na2SO4; 50 µM FeCl3 (sterilized by filtration at 0.22 µm); 2 µM MnSO4; 8 g l^-1^ glucose; 2 g l^-1^ L-glutamic acid; pH 7.0), 1 µg of DNA recombinant fragment was added to the GA1 cells suspension adjusted to an OD600_nm_ of 0.01 into MMG liquid medium. One day after incubation at 37 °C with shaking at 165 rpm, bacteria were spread on LB plates supplemented with the appropriate antibiotic to select positive colonies.

Construction of deletion mutants of *Pseudomonas* sp. CMR12a

E-PCH and PVD mutants of *Pseudomonas* sp. CMR12a were constructed using the I-SceI system and the pEMG suicid vector^63, 64^. Briefly, the upstream and downstream region flanking the *pchA* (*C4K39_5481*) or the *pvdI* (*C4K39_6027*) genes were PCR amplified (primers listed in the Supplementary Table 3), linked via overlap PCR and inserted into the pEMG vector. The resulting plasmid (Supplementary Table 3) was integrated by conjugation into the *Pseudomonas* sp. CMR12a chromosome via homologous recombination. Kanamycin (25µg/mL) resistant cells were selected on King B agar plates and transformed by electroporation with the pSW-2 plasmid (harboring I-SceI system). Gentamycin (20µg/ml) resistant colonies on agar plates were transferred to King B medium with and without kanamycin to verify the loss of the antibiotic (kanamycin) resistance. *Pseudomonas* mutants were identified by PCR with the corresponding UpF and DwR specific primers and via the loss of E-PCH or/and PVD production (see section Secondary metabolites analysis).

### *Pseudomonas* sp. cell-free supernatant

*Pseudomonas* sp. strains were grown overnight on LB solid medium, at 30 °C. The cell suspension was adjusted to OD600nm 0.05 by resuspension in 100 ml of CAA and when appropriate supplemented with 20 µg/l of FeCl3.6H2O (iron supplementation). Cultures were shaken at 120 rpm at 30 °C for 48 h and then centrifuged at 5000 rpm at room temperature (22 °C) for 20 min. The supernatant was filter-sterilized (0.22 µm pore size filters) and stored at -20 °C until use.

### Dual interactions

*B. velezensis* strains were grown overnight on LB solid medium, at 30 °C. Cells were resuspended in 2 ml of EM liquid medium to a final OD600nm of 0.1 in which 1, 2, or 4% v/v (depending on the experiment and indicated in the figures legends) of *Pseudomonas* CFS were added while the control remained un-supplemented. *B. velezensis* liquid cultures were shaken in an incubator at 300 rpm at 30 °C for 24 h. Additionally, 2 ml of the (co-)culture supernatants were sampled at 8 h and 24 h, centrifugated at 5000 rpm at room temperature (approx. 22 °C) for 10 min to extract supernatants and collect the cells. Further, cell-free (co-)culture supernatants were filter-sterilized (0.22µm) and used for analytical analysis of secondary metabolites and antibacterial assays. For some experiments using 2H-bae as a marker, the CFS obtained from the double mutant sessilins and orfamides (Δ*sesA-ofaBC*) was used instead of CFS from CMR12a wild-type because it yielded a higher response and lower inhibition interferences by CLPs. The remaining cells, after supernatant collection, were stored at -80 °C to avoid RNA degradation, until performing RT-qPCR analysis.

### Antimicrobial activity assays

Antibacterial activity of the *B. velezensis* supernatant generated after dual interaction with *Pseudomonas* CFS was tested against *X. campestris* pv. *campestris* and *C. michiganensis* subsp. *michiganensis.* The activity of co-culture supernatants was quantified in microtiter plates (96-well) filled with 250 µl of LB liquid medium, inoculated at OD_600nm_ = 0.1 with *X. campestris* pv. *Campestris* and *C. michiganensis* subsp. *michiganensis* and supplemented with 2% or 6% v/v of the supernatants, respectively. The activity of (co-)culture supernatants was estimated by measuring the pathogen OD_600nm_ every 30min during 24 h with a Spectramax® (Molecular Devices, Wokingham, UK), continuously shaken, at 30 °C. For estimating the activity of co-culture supernatants on a solid medium, 5 µl supernatant was applied to a sterile paper disk (5 mm diameter). After drying, disks were placed on LBA square plates previously inoculated with a confluent layer of *X. campestris* pv. *campestris*, *C. michiganensis* subsp. *michiganensis, P. carotovorum, P. fuscovaginae, P. cichorii, A. tumefaciens or R. fascians*. LB liquid medium was used as a negative control. Plates were incubated at 25 °C for 48 h. Three repetitions were done and the inhibition zones from the edge of the paper discs to the edge of the zone were measured. Antibacterial activity of the different *Pseudomonas* strains on the *B. velezensis* strains growth were tested by adding different % (v/v) of the corresponding *Pseudomonas* CFS in microtiter plates (96-well) filled with 250 µl of EM liquid medium. *B. velezensis* OD600nm after 7 h was measured with a Spectramax® (Molecular Devices, Wokingham, UK).

### RNA isolation and RT-qPCR

RNA extraction and DNAse treatment were carried out using the NucleoSpin RNA Kit (Macherey Nagel, Germany), following the Gram + manufacturer’s protocol. RNA quality and quantity were performed with Thermo scientific NanoDrop 2000 UV-vis Spectrophotometer. Primer 3 program available online was used for primer design and primers were synthesized by Eurogentec. The primer efficiency was evaluated and primer pairs showing an efficiency between 90 and 110% in the qPCR analysis were selected. Reverse transcriptase and RT-qPCR reactions were conducted using the Luna® Universal One-Step RT-qPCR Kit (New England Biolabs, Ipswich, MA, United States). The reaction was performed with 50 ng of total RNA in a total volume of 20 µL: 10 µL of luna universal reaction mix, 0.8 µL of each primer (10 µM), 5 µL of cDNA (50ng), 1 µL of RT Enzyme MIX, 2.4 µl of Nuclease-free water. The thermal cycling program applied on the ABI StepOne was: 55 °C for 10 min, 95 °C for 1 min, 40 cycles of 95 °C for 10 s and 60 °C for 1 min, followed by a melting curve analysis performed using the default program of the ABI StepOne qPCR machine (Applied Biosystems). The real-time PCR amplification was run on the ABI step-one qPCR instrument (Applied Biosystems) with software version 2.3. The relative gene expression analysis was conducted by using the 2*ΔCt* method^65^ with the *gyrA* gene as a housekeeping gene to normalize mRNA levels between different samples. The target genes in this study were *dfnA*, *baeJ* and *acnA*.

### Secondary metabolite analysis

For detection of BSMs, *B. velezensis* and *Pseudomonas* sp. were cultured in EM and CAA as described above. After an incubation period of 24 h for *B. velezensis*, if not differentially indicated, and 48 h for *Pseudomonas* sp., supernatants of the bacteria were collected and analyzed by UPLC MS and UPLC qTOF MS/MS. Metabolites were identified using Agilent 1290 Infinity II coupled with DAD detector and Mass detector (Jet Stream ESI-Q-TOF 6530) in both negative and positive mode with the parameter set up as follows: parameters: capillary voltage: 3.5 kV; nebulizer pressure: 35 psi; drying gas: 8 l/min; drying gas temperature: 300 °C; flow rate of sheath gas: 11 l/min; sheath gas temperature: 350 °C; fragmentor voltage: 175 V; skimmer voltage: 65 V; octopole RF: 750 V. Accurate mass spectra were recorded in the range of m/z = 40-250. An C18 Acquity UPLC BEH column (2.1 × 50 mm × 1.7 µm; Waters, milford, MA, USA) was used at a flow rate of 0.3 ml/min and a temperature of 40 °C. The injection volume was 20 µl and the diode array detector (DAD) scanned a wavelength spectrum between 190 and 600 nm. A gradient of 0.1% formic acid water (solvent A) and acetonitrile acidified with 0.1% formic acid (solvent B) was used as a mobile phase with a constant flow rate at 0.45 ml/min starting at 10% B and raising to 100% B in 20 min. Solvent B was kept at 100% for 2 min before going back to the initial ratio. Secondary metabolite quantification was performed by using UPLC–MS with UPLC (Acquity H-class, Waters) coupled to a single quadrupole mass spectrometer (SQD mass analyzer, Waters) using a C18 column (Acquity UPLC BEH C18 2.1 mm × 50 mm, 1.7 µm). Elution was performed at 40 °C with a constant flow rate of 0.6 ml/min using a gradient of Acetonitrile (solvent B) and water (solvent A) both acidified with 0.1% formic acid as follows: 2 min at 15% B followed by a gradient from 15% to 95% during 5 min and maintained at 95% up to 9.5 min before going back to initial conditions at 10 min during 2 min before next injection. Compounds were detected in both electrospray positive and negative ion mode by setting SQD parameters as follows: cone voltage: 60V; source temperature 130 °C; desolvation temperature 400 °C, and nitrogen flow: 1000 l/h with a mass range from m/z 300 to 2048. 3D chromatograms were generated using the open-source software MzMine 2^66^.

### Bioguided fractionation

*Pseudomonas* CFS were concentrated with a C18 cartridge ‘Chromafix, small’ (Macherey-Nagel, Düren, Germany). The column was conditioned with 10 ml of MeOH followed by 10 ml of milliQ water. Then, 20 ml of supernatant flowed through the column. The metabolites were eluted with 1 ml of a solution of increasing acetonitrile/water ratio from 5:95 to 100:0 (v/v). The triggering effect of these fractions on *Bacillus* 2H-bae production was tested in 48 wells microplate containing 1 ml of EM medium inoculated with *B. velezensis* GA1 (OD_600nm_ = 0.1) and 4% v/v of aforementioned *Pseudomonas* fractions, growing for 24 h, with shaking at 300 rpm and 30 °C. Afterward, the production of 2H-bae was quantified compared to controls and crude supernatant.

### Purification of E-PCH and PVD

PVD and E-PCH were purified in two steps. Firstly, *Pseudomonas* CFS were concentrated with a C18 cartridge (as indicated in section Bioguided fractionation) and eluted with 2 times 2 ml of a solution of water and ACN (15 and 30% of ACN (v/v)). Secondly, the fractions were injected on HPLC for purification performed on an Eclipse+ C18 column (L = 150 mm, D = 3.0 mm, Particles diameter 5 µm) (Agilent, Waldbronn, Germany). The volume injected was 100 µl. The UV-Vis absorbance was measured with a VWD Agilent technologies 1100 series (G1314A) detector (Agilent, Waldbronn, Germany). The lamp used was a Deuterium lamp G1314 Var Wavelength Det. (Agilent, Waldbronn, Germany). Two wavelengths were selected: 320 nm, used for the detection of E-PCH, and 380 nm, used for the detection of PVD. The fractions containing the PVD and E-PCH were collected directly at the detector output. Further, the purity of the samples was verified by two detectors, a diode array detector (DAD) 190 to 601 nm (steps: 1 nm) and a Q-TOF (tandem mass spectrometry, quadrupole and Time of flight detector combined) (Agilent, Waldbronn, Germany). Electrospray ionization was performed in positive mode (ESI+) (Dual AJS ESI) (Vcap = 3500 V, Nozzle Voltage = 1000 V), with a mass range from m/z 200 to 1500. Finally, the concentration of PVD and E-PCH were estimated by utilization of Beer-Lambert law formula, A = Ɛlc (A: absorbance; Ɛ: molar attenuation coefficient or absorptivity of the attenuating species; l: optical path length and c: concentration of molecule). l value for E-PCH and PVD is 1 cm while Ɛ is 4000 L.mol^-1^.cm^-1^ or 16000 L.mol^-1^.cm^-1^, respectively^67^. The absorbance was measured with VWR, V-1200 Spectrophotometer, at 320 nm (pH = 8) for E-PCH and 380nm (pH = 5) for PVD^67^. Further, the absorbance value was used for calculating the final concentration. The fragmentation pattern of *Pseudomonas* sp. CMR12a PVD was obtained by UPLC MS/MS analysis of m/z = 1288.5913 ion in positive mode with fragmentation energy at 75 V and compared to the one described in *P. protegens* Pf-5^40^.

### Confrontation, white line formation and motility test

For confrontation assays on agar plates*, Bacillus* and *Pseudomonas* strains were grown overnight in EM and CAA liquid mediums, respectively. After bacterial washing in peptone water and adjustment of OD_600nm_ to 0.1, 5 µl of bacterial suspension was spotted at 1 mm, 5 mm and 7.5 mm distance onto an EM agar plate. For the white line formation experiments, *B. velezensis* line was applied with a cotton stick and 5 µl of *Pseudomonas* sp. cell suspensions were spotted at a 5 mm distance onto CAA agar plates. Plates were incubated at 30 °C and images taken after 24 h.

Photographs were captured using CoolPix camera (NiiKKOR 60x WIDE OPTICAL ZOOM EDVR 4.3-258 mm 1:33-6.5).

### MALDI-FT-ICR MS imaging

Mass spectrometry images were obtained as recently described^68^ using a FT-ICR mass spectrometer (SolariX XR 9.4T, (Bruker Daltonics, Bremen, Germany)) mass calibrated from 200 m/z to 2,300 m/z to reach a mass accuracy of 0.5 ppm. Region of interest from agar microbial colonies was directly collected from the Petri dish and transferred onto an ITO Glass slide (Bruker, Bremen, Germany), previously covered with double-sided conductive carbon tape. The samples were dried under vacuum and covered with an α-cyano-4-hydroxycinnamic acid (HCCA) matrix solution at 5 mg/mL (70 : 30 acetonitrile : water v/v). In total, 60 layers of HCCA matrix were sprayed using the SunCollect instrument (SunChrom, Friedrichsdorf, Germany). FlexImaging 5.0 (Bruker Daltonics, Bremen, Germany) software was used for MALDI-FT-ICR MS imaging acquisition, with a pixel step size for the surface raster set to 100 µm.

### *In planta* competition

For *in planta* studies, tomato seeds (*Solanum lycopersicum* var. Moneymaker) were sterilized in 75% ethanol with shaking for 2 min. Subsequently, ethanol was removed and seeds were added to the 50 ml sterilization solution (8.5 ml of 15% bleach, 0.01 g of Tween 80 and 41.5 ml of sterile ultra-pure water) and shaken for 10 min. Seeds were thereafter washed five times with water to eliminate stock solution residues. Further, seeds were placed on square Petri dishes (5 seeds/plate) containing Hoagland solid medium (14 g/l agar, 5 ml stock 1 (EDTA 5,20 mg/l; FeSO_4_x7H_2_O 3,90 mg/l; H_3_B0_3_ 1,40 mg/l; MgSO_4_x7H_2_O 513 mg/l; MnCl_2_x4H_2_O 0,90 mg/l, ZnSO_4_x7H_2_O 0,10 mg/l; CuSO_4_x5H_2_O 0,05 mg/l; 1 ml in 50 ml stock 1, NaMo0_4_x2H_2_O 0,02 mg/l 1 ml in 50 ml stock 1), 5 ml stock 2 (KH_2_PO_4_ 170 mg/l; 5 ml stock 3: KN0_3_ 316 mg/l, Ca(NO_3_)_2_ 4H_2_O 825 mg/l), pH = 6,5) and placed in the dark for three days. Afterwards, 10 seeds were inoculated with 2 µl of overnight culture (OD_600_ = 0.1) of the appropriate strains (control) or with a mix of *Bacillus* and *Pseudomonas* cells (95:5 ration) (interaction) and grown at 22 °C under a 16/8 h night/day cycle with constant light for three days. After the incubation period, to determine bacterial colonization levels, bacteria from roots of six plants per condition were detached from roots by vortexing for 1 min in peptone water solution supplemented with 0.1% of Tween. Serial dilutions were prepared and 200 µl of each were plated onto LB medium using plating beads. After 24 h of incubation at 30 °C for *Pseudomonas* and at 42 °C for *Bacillus*, colonies were counted. Colonization results (six plants per strain) were log-transformed and statistically analyzed. Three independent assays were performed with six plants each for *in planta* competition assays. To measure bacterial BSMs production *in planta*, a rectangle part (1 x 2.5 cm) of medium close to the tomato roots was sampled. BSMs were extracted for 15 min, with 1.5 ml of acetonitrile (85%). After centrifugation for 5 min at 4000 rpm, the supernatant was recovered for UPLC-MS analysis as previously described.

### Statistical analysis

Statistical analyses were performed using GraphPad PRISM software with Student paired T-test or Mann-Whitney test. For multiple comparisons, one-way ANOVA and Tukey tests were used in RStudio 1.1.423 statistical software environment (R language version 4.03)^69^.

## Acknowledgments

We gratefully acknowledge Sébastien Rigali, Alexandre Jousset and Loïc Ongena for critically reading the manuscript. We thank C. Keel for the kind gift of strains and J. Vacheron for the very helpful indications on *Pseudomonas* mutagenesis. This work was supported by the EU Interreg V France-Wallonie-Vlaanderen portfolio SmartBiocontrol (Bioprotect and Bioscreen projects, avec le soutien du Fonds européen de développement régional - Met steun van het Europees Fonds voor Regionale Ontwikkeling), by the European Union Horizon 2020 research and innovation program under grant agreement No. 731077 and by the EOS project ID 30650620 from the FWO/F.R.S.- FNRS. The MALDI FT-ICR SolariX XR was funded by FEDER BIOMED HUB Technology Support (number 2.2.1/996). AA is recipient of a F.R.I.A. fellowship (Formation à la Recherche dans l’Industrie et l’Agriculture) and MO is senior research associate at the F.R.S.-F.N.R.S.

## Author contributions

SA, TM, AR and AA performed most of the co-culture and *in planta* experiments. SA and TM performed most of molecular biology experiments with help of GH and SS for mutant generation and of SS for transcriptomics. TM and GH did genome mining. AR and AA were involved in all aspects of metabolomics using UPLC-MS. Data analysis was done by SA, TM, AA and AR. AM, AA and EDP performed the MALDI FT-ICR experiments and analyzed the data. MH and RDM provided *Pseudomonas* strains/mutants and also supported the study by providing intellectual input. SA, TM and MO mainly wrote the manuscript. All of the authors commented on the manuscript and contributed to the final form. MO supervised the study.

