## Supplementary material for "Chelator sensing and lipopeptide interplay mediates molecular interspecies interactions between soil bacilli and pseudomonads": Suppl tables

**Supplementary Table 1. Prediction and detection of *B. velezensis* GA1 and *Pseudomonas* sp. CMR12a secondary metabolites.** Biosynthetic gene clusters were predicted by antiSMASH 5.0[21] and the corresponding metabolites were detected (+) or not (ND) by our UPLC-qTOF-MS/MS method. The locus tag numbers for *Pseudomonas* sp. CMR12a corresponds to the genome annotation (Genbank: CP027706.1). The locus tag for *B. velezensis* GA1 corresponds to the recently sequenced genome annotation (Genbank: CP046386.1).

| BSM predicted | Category | Genes involved in the biosynthesis | Disrupted gene in the non-producing mutant | Detected |
|---|---|---|---|---|
| ***B. velezensis* GA1** | | | | |
| Surfactins | NRPs/CLPs | *srfaABCD* | Δ*srfaA* | + |
| Fengycins | NRPs/CLPs | *fenABCDE* | Δ*fenA* | + |
| Iturins | NRPs/CLPs | *ituDABC* | Δ*ituA* | + |
| Macrolactins | NRPs/PKs | *mlnABCDEFGHI* | Δ*mlnA* | + |
| Bacillaene | NRPs/PK | *baeBCDEGHIJLMNRS* | Δ*baeS* and Δ*baeJ* | + |
| Dihydrobacillaene | NRPs/PK | *baeBCDEGHIJLMNRS* | Δ*baeJ* | + |
| Difficidin | NRPs/PK | *loaP-dfnABDEFGHIJKLMNO* | Δ*dfnM* | + |
| Oxydifficidin | NRPs/PK | *loaP-dfnABDEFGHIJKLMNO* | Δ*dfnM* and Δ*dfnA* | + |
| Bacillibactin | NRPs/Siderophore | *dhbACEBF* | Δ*dhbC* | + |
| Bacilysin | Dipeptide | *bacABC* | Δ*bacA* | ND |
| Amylocyclicin | RiPP | *acnBACDEF* | Δ*acnA* | ND |
| Amylolysin | RiPP | *amlFEKRAMT* | Δ*amlA* | ND |
| ***Pseudomonas* sp. CMR12a** | | | | |
| Sessilins | NRPs/CLPs | *sesTRDABC-macA1B1* | Δ*sesA* | + |
| Orfamides | NRPs/CLPs | *ofaR1ABC-macA2B2-ofaR2* | Δ*ofaBC* | + |
| Phenazines | Antibiotics | *phzABCDEFGH* | Δ*phzABCDEFGH* | + |
| Pyoverdine | NRPs/Siderophore | *pvdYSLH-C4K39_6099* to *C4K39_6098-fpvCDEF-C4K39_6092* to *C4K39_6089-C4K39_6029* to *C4K39_6028-pvdIJD-fpvZ-pvdEFONMP-ampO-pvdTR-fvpI-pvdA* | Δ*pvdL* | + |
| Enantio-pyochelin | NRPs/Siderophore | *pchR-pchDHIEFKCBA* | Δ*pchA* | + |

**Supplementary Table 2: Conservation of genes encoding substrate binding proteins involved in iron transport in *B. subtilis* and *B. velezensis***. The percentage of genes nucleotide identity was obtained by blast comparison performed on the MAGE platform (https://mage.genoscope.cns.fr).

| | ***B. velezensis* GA1** | | ***B. velezensis* QST713** | | ***B. velezensis* S499** | | ***B. velezensis* FZB42** | |
|---|---|---|---|---|---|---|---|---|
| **Transporters in *B. subtilis* 168** | **Gene name** | **% of identity** | **Gene name** | **% of identity** | **Gene name** | **% of identity** | **Gene name** | **% of identity** |
| *feuA* | *GL331_06035* | 83 | *BVQ_00990* | 83 | AS588_13145 | 83 | RBAM_002120 | 83 |
| *fhuD* | *GL331_01180* | 81 | *BVQ_17365* | 82 | AS588_15255 | 82 | RBAM_030440 | 82 |
| *yxeB (frxB)* | *GL331_04235* | 72 | *BVQ_20555* | 73 | AS588_18310 | 72 | RBAM_036560 | 72 |
| *fpiA* (*pbtQ/yclQ)* | *GL331_06955* | 74 | *BVQ_01985* | 73 | AS588_12230 | 74 | RBAM_004080 | 73 |
| *yfmC (fecC)* | *GL331_10185* | 36 | *BVQ_05570* | 36 | AS588_10320 | 36 | RBAM_010510 | 37 |
| *yfiY(sxzY)* | *GL331_04235 (yxeB)* | 33 | *BVQ_20555 (yxeB)* | 32 | AS588_18310 (*yxeB*) | 33 | RBAM_036560 (*yxeB*) | 32 |

**Supplementary Table 3. Strains and plasmids used in this study.**

| | Relevant genotype and description | References or sources |
|---|---|---|
| ***Bacillus velezensis*** | | |
| GA1 | Wild type | [1] |
| GA1 Δ*sfp* ::cat | GA1 deleted of *sfp* gene; unable to produce lipopeptides, polyketides and bacillibactin | This study |
| GA1 Δ*srfaA* ::cat | GA1 deleted of *srfaA* gene; unable to produce surfactins | This study |
| GA1 Δ*ituA* ::cat | GA1 deleted of *ituA* gene; unable to produce iturins | This study |
| GA1 Δ*fenA* ::cat | GA1 deleted of *fenA* gene; unable to produce fengycins | This study |
| GA1 Δ*fenA* ::cat-Δ*ituA* ::phleo-Δ*srfaA* ::kan | GA1 deleted of *fenA*, *ituA* and *srfaA* genes; unable to produce surfactins, fengycins and iturins | This study |
| GA1 Δ*baeJ* ::cat | GA1 deleted of *baeJ* gene; unable to produce bacillaene and dihydrobacillaene | This study |
| GA1 Δ*baeS* ::cat | GA1 deleted of *baeS* gene; unable to produce bacillaene | This study |
| GA1 Δ*dfnA* ::cat | GA1 deleted of *dfnA* gene; unable to produce difficidin and oxydifficidin | This study |
| GA1 Δ*dfnM* ::cat | GA1 deleted of *dfnM* gene; unable to produce oxydifficidin | This study |
| GA1 Δ*baeJ* ::cat Δ*dfnA* ::phleo | GA1 deleted of *baeJ* and *dfnM* genes; unable to produce bacillaenes and difficidins | This study |
| GA1 Δ*mlnA* ::cat | GA1 deleted of *mlnA* gene; unable to produce macrolactins | This study |
| GA1 Δ*dhbC* ::cat | GA1 deleted of *dhbC* gene; unable to produce bacillibactin | This study |
| GA1 Δ*acnA* ::cat | GA1 deleted of *acnA* gene; unable to produce amylocyclicin | This study |
| GA1 Δ*acnA*::cat Δ*sfp*::phleo | GA1 deleted of *sfp* and *acnA* genes; unable to produce lipopeptides, polyketides, bacillibactin and amylocyclicin | This study |
| GA1 Δ*amlA* ::cat | GA1 deleted of *amlA* gene; unable to produce amylolysin | This study |
| GA1 Δ*amlA*::cat Δ*sfp*::phleo | GA1 deleted of *sfp* and *amlA* gene; unable to produce lipopeptides, polyketides, bacillibactin and amylolysin | This study |
| GA1 Δ*bacA* ::cat | GA1 deleted of *bacA* gene; unable to produce bacilysin | This study |
| GA1 Δ*bacA*::cat Δ*sfp*::phleo | GA1 deleted of *sfp* and *bacA* gene; unable to produce lipopeptides, polyketides, bacillibactin and bacylisin | This study |

| | | |
|---|---|---|
| ***Pseudomonas* sp.** | | |
| CMR12a | Wild type | [2] |
| Δ*sesA* | CMR12a disrupted of *sesA* gene; $Gm^R$; unable to produce sessilins | [3] |
| Δ*ofaBC* | CMR12a deleted of *ofaB* and *ofaC* genes; $Gm^R$; unable to produce orfamides | [4] |
| Δ*phz* | CMR12a deleted of phenazine biosynthesis operon; unable to produce phenazines | [3] |
| Δ*sesA-ofaBC* | CMR12a disrupted of *sesA* gene and deleted *ofaB* and *ofaC* genes; $Gm^R$; unable to produce sessilins and orfamides | [4] |
| Δ*sesA-phz* | CMR12a disrupted of *sesA* gene and deleted phenazines biosynthesis operons; $Gm^R$; unable to produce sessilins and phenazines | [3] |
| Δ*ofaAC-phz* | CMR12a deleted of *ofaB* and *ofaC* genes and phenazines biosynthesis operons; unable to produce orfamides and phenazines | [4] |
| Δ*sesA-ofaBC-phz* | CMR12a disrupted of *sesA* gene and deleted *ofaB* and *ofaC* genes and phenazine biosynthesis operons; $Gm^R$; unable to produce sessilins, orfamides and phenazines | [4] |
| Δ*pchA* | CMR12a deleted of *pchA* gene; unable to produce enantio-pyochelin | This study |
| Δ*pvdI* | CMR12a deleted of *pvdI* gene; unable to produce pyoverdine | This study |
| Δ*pvdI-pchA* | CMR12a deleted of *pvdI* and *pchA* genes; unable to produce pyoverdine and enantio-pyochelin | This study |
| ***Pseudomonas putida*** | | |
| RW10S2 | Wild type; WLIP producer | [5] |
| BW11M1 | Wild type; xantholysins producer | [6] |
| WCU-64 | Wild type; putisolvins producer | [7] |
| ***Pseudomonas lactis*** | | |
| SS101 | Wild type; massetolides producer | [8] |
| ***Pseudomonas tolaasii*** | | |
| CH36 | Wild type; tolaasins and pseudodesmin producer | [5] |
| CH36Δ*tolA* | CH36 disrupted of *tolA* gene; $Gm^R$; unable to produce tolaasins | [5] |
| ***E. coli*** | | |

| | | |
|---|---|---|
| *DH5αpir* | supE44, ΔlacU169 (ΦlacZΔM15), recA1, endA1, hsdR17, thi-1, gyrA96, relA1, λpir | 9 |
| *DH5α p497* | Helper strain harboring P497 plasmid | C. Keel laboratory |
| **Plasmids** | | |
| pEMG | pSEVA212S; oriR6K, *lacZα* with two flanking I-SceI sites; $Km^r$, $Ap^r$ | 10 |
| pEMG-*pchA* | Suicide plasmid used for the deletion of *pchA* | This study |
| pEMG-*pvdI* | Suicide plasmid used for the deletion of *pvdI* | This study |
| pSW-2 | oriRK2, *xylS*, $P_m$::*I-sceI*; $Gm^R$ | 10 |
| **Phytopathogenic strains** | | |
| *Xanthomonas campestris* pv. *campestris* | | DSMZ¹ N°3586 |
| *Clavibacter michiganensis* subsp. *michiganensis* | | DSMZ¹ N°20741 |
| *Pectobacterium carotovorum* | | De Mot laboratory |
| *Pseudomonas fuscovaginae* | | De Mot laboratory |
| *Pseudomonas cichorii* | | De Mot laboratory |
| *Agrobacterium tumefaciens* | | De Mot laboratory |
| *Rhodoccocus fascians* | | De Mot laboratory |

### Supplementary Table 4. Primers used in this study

| | Primer Name | Primer sequence (5'->3') | Targeted genes |
|---|---|---|---|
| **Deletion mutant** | | | |
| ***B. velezensis* GA1** | | | |
| | UpsrfaAF | TCAGCAAAACTGCGTGGTAG | *srfaA* |
| | UpsrfaAR | CCAATTTTCGAATTCTTTTACCGCGATAAAAAGTTATTTCCATATGTGTGC | |
| | DwsrfAF | CAGCTCCAGATCCTCTACGCCGGACACGCTTTATATCGTGCCGAA | |
| | DwsrfAR | AAGAAATGATCATAAATACC | |
| | UpFenAF | AGCAAAAACCGGGTCACTAA | *fenA* |
| | UpFenAR | CCAATTTTCGAATTCTTTTACCGCGTTCGTCTGACATGACAAGCA | |
| | DwFenAF | CAGCTCCAGATCCTCTACGCCGGACAAAGGACTTTAATTTCATAAAAAGGTG | |
| | DwFenAR | CCTTTTTGAGAAGAGAAGAAAAAG | |
| | UpItuAF | ATGCAGGAAATAGGGGTGAA | *ituA* |
| | UpItuAR | CCAATTTTCGAATTCTTTTACCGCGGGTATACATAGGTCCCCTCCTG | |
| | DwItuAF | CAGCTCCAGATCCTCTACGCCGGACCAATTGAACTTTTAGGGAAAAGCA | |
| | DwItuAR | GCGACTAACGTATCGGGTTG | |
| | UpDfnAF | GACTTTTGAATAATCTACAGTGTCTCC | *dfnA* |
| | UpDfnAR | TTTTCGAATTCTTTTACCGCGAAACGCGTTTGCGATTCAG | |
| | UpDfnAphleoR | CAGGAAACAGCTATGACAAACGCGTTTGCGATTCAG | |
| | DwDfnAphleoF | GTAAAACGACGGCCAGTACAGGCTGAGTATGACCAGACA | |
| | DwDfAF | CAGCTCCAGATCCTCTACGCCGGACACAGGCTGAGTATGACCAGACA | |
| | DwDfnAR | TCCGGAATATGATCTTGTGAAG | |
| | UpDfnMF | GGGCAGTGGAGCTGTACC | *dfnM* |
| | UpDfnMR | CCAATTTTCGAATTCTTTTACCGCGGGTCATTTTCATTCCTCCAAGA | |
| | DwDfnMF | CAGCTCCAGATCCTCTACGCCGGACCTTGTTGAGTTTTGAACGAAAAA | |
| | DwDfnMR | AGCCGTTATCAATCGTGCTG | |
| | UpBaeJF | GTATGCGTCCCAGACTCAGC | *baeJ* |
| | UpBaeJR | CCAATTTTCGAATTCTTTTACCGCGTTTCATAGAGCTGCCTCCAT | |
| | DwBaeJF | CAGCTCCAGATCCTCTACGCCGGACGGGATACCTATGAAGTGGAGGTT | |
| | DwBaeJR | TCATAGTAGCCGACTTGAGAATCA | |
| | UpBaeSF | GTACAGCAAGGTGCCATGAG | *baeS* |
| | UpBaeSR | CCAATTTTCGAATTCTTTTACCGCGTTTTTGAAAAGACATAACCAACAG | |
| | DwBaeSF | CAGCTCCAGATCCTCTACGCCGGACTTTTAATATCGCCCCCTGTTT | |
| | DwBaeSR | GAGGCGTTGAAGCATACCAG | |
| | UpsfpF | TCGTCACCCATGAAATCAAA | *sfp* |
| | UpsfpR | CCAATTTTCGAATTCTTTTACCGCGCATGTCCAGATCCTCCGTCT | |
| | DwsfpF | CAGCTCCAGATCCTCTACGCCGGACGACGGGATTGAGATGAAAA | |
| | DwsfpR | CATTGAGACGTACCCGCTTT | |
| | UpdhbCF | GCGTTTCTGCCTGAATCC | *dhbC* |
| | UpdhbCR | CCAATTTTCGAATTCTTTTACCGCGCATGTTTGTCCCTCCTTTTCGT | |
| | DwdhbCF | CAGCTCCAGATCCTCTACGCCGGACGGCTTTACCAAGATGA | |
| | DwdhbCR | GCAGCACTTGAAGGCTTGAT | |
| | UpmlnAF | CGGAAAAACCGTTTCAAAAA | *mlnA* |
| | UpmlnAR | CAGGAAACAGCTATGACTTTTAAAATTGTCATTTACTCTAAGCA | |
| | DwmlnAF | GTAAAACGACGGCCAGTCTAAGGCGCAGATTGGATA | |
| | DwmlnAR | TGTACCTGTGCCATGTGCTT | |
| | UpbacAF | GATGGGTCTGATCGTGTCAA | *bacA* |
| | UpbacAR | CCAATTTTCGAATTCTTTTACCGCGCATGAGCACCAACCAATCTG | |
| | DwbacAF | CAGCTCCAGATCCTCTACGCCGGACAACTGAACAAGATTTGCAGG | |
| | DwbacAR | GAATCGGGGCGACAATTT | |
| | UpacnAF | TCCTTGTCACTGGGTGATGA | *acnA* |
| | UpacnAR | TTTCGAATTCTTTTACCGCGGTTCATCATAACATCTCCCTACTCTG | |
| | DwacnAF | CCAGATCCTCTACGCCGGACGCAGCTGCTTGGTAAAATCG | |
| | DwacnAR | CGCAAAATCAGCGTTTGTC | |

| | | | |
|---|---|---|---|
| | UpamlAF | GGGCTGACAGGGATAAAAGA | *amlA* |
| | UpamlAR | TTTCGAATTCTTTTACCGCGCTCATTCATTAATATTCCTCCCTTTG | |
| | DwamlAF | CCAGATCCTCTACGCCGGACTGGTGTTAAAACAACCCGAAA | |
| | DwamlAR | TCATGATCTCTAATTTTCTCATTCAAA | |
| ***Pseudomonas* sp. CMR12a** | | | |
| | UppvdIF | GGCATTCTTGACCGGTCGTC | *pvdI* |
| | UppvdIR | GTGTTGTCCATTACACAGCCTCCATTGCATTCATCGGGAGTCATCC | |
| | DwpvdIF | ATGGAGGCTGTGTAATGGACAACA | |
| | DwpvdIR | TGTAGCGGTGTAGCAGAG | |
| | pvdICheckF | CCTGCTGCTGGAAGGATTGA | |
| | pvdICheckR | GGATCGAGCTGCCAAAGGAA | |
| | UppchAF | GACCAACTGCCGGCGGAT | *pchA* |
| | UppchAR | CCTTCAGCGATCGGCCGGTGCATCACATCTTGCGCTCCTTGCTCC | |
| | DwpchAF | TGATGCACCGGCCGATC | |
| | DwpchAR | GTGGTGAAGCTTTCCATGCC | |
| | pchACheckF | TCATCCACTGGAACATCGCC | |
| | pchACheckR | GCGGACTGATTTCCTCGGTA | |
| **Antibiotic marker** | | | |
| | CatF | CGCGGTAAAAGAATTCGAAAA | Chloramphenicol marker |
| | CatR | GTCCGGCGTAGAGGATCTG | |
| | PhleoF | GTCATAGCTGTTTCCTGCCAAAAGGGGGTTTCATTTT | Phleomycin marker |
| | PhleoR | ACTGGCCGTCGTTTTACTCCAATAAATGCGACACCAA | |
| | KanF | GTCATAGCTGTTTCCTGCTTAGCTCCTGAAAATCTCGG | Kanamycin marker |
| | KanR | ACTGGCCGTCGTTTTACCTGATAATTACTAATACTAGGAGAAG | |
| | nptIIF | GAGGATCGTTTCGCATGATT | Kanamycin marker for *Pseudomonas* |
| | nptIIR | CGCTCAGAAGAACTCGTCAA | |
| | psw-F | GGACGCTTCGCTGAAAACTA | pSW-II insertion |
| | psw-R | AACGTCGTGACTGGGAAAAC | |
| **RT-qPCR** | | | |
| ***B. velezensis* GA1** | | | |
| | AcnA_F_qPCR | CCAAGCAGCTGCGTATTTTT | *acnA* |
| | AcnA_R_qPCR | CTTCGACTCTGGGCATCTCT | |
| | QgyrA_F | GAGACGCACTGAAATCGTGA | *gyrA* |
| | QgyrA_R | GCCGGGAGACGTTTAACATA | |
| | BaeJQ_F_qPCR | CCGATGACGATTCCTGAAGT | *baeJ* |
| | BaeJQ_R_qPCR | GCCCTTTCACAATCGAAAGA | |
| | DfnA_F_qPCR | GGCGTTTTTGCCTCTTCGTT | *dfnA* |
| | DfnA_R_qPCR | ATCAGACGGCGTATCGTGTC | |
