## Supplementary material for "Chelator sensing and lipopeptide interplay mediates molecular interspecies interactions between soil bacilli and pseudomonads": Suppl figures

**a**

| Bacillus velezensis GA1 and Pseudomonas sp. CMR12a detectable Bioactive Secondary Metabolite (BSMs) differential production | | | | | |
|---|---|---|---|---|---|
| BSM | Main described activities | Raw formula of variants | Detected m/z (mass error (ppm)) | EM | CAA |
| 1 Surfactins | ISR Motility | **C12** : $C_{50}H_{87}N_7O_{13}$<br>**C13** : $C_{51}H_{89}N_7O_{13}$<br>**C14** : $C_{52}H_{91}N_7O_{13}$<br>**C15** : $C_{53}H_{93}N_7O_{13}$<br>**C16** : $C_{54}H_{95}N_7O_{13}$ | 994.6420 (1,5)<br>1008.6582 (0,9)<br>1022,6743 (0,5)<br>1036.6904 (4,8)<br>1050.7061 (1,7) | | |
| 2 Iturins | Antifungal | **C14** : $C_{48}H_{74}N_{12}O_{14}$<br>**C15** : $C_{49}H_{76}N_{12}O_{14}$<br>**C16** : $C_{50}H_{78}N_{12}O_{14}$<br>**C17** : $C_{51}H_{80}N_{12}O_{14}$ | 1043.5510 (1,0)<br>1057.5653 (2,2)<br>1071.5809 (2,3)<br>1085.5967 (2,1) | | |
| 3 Fengycins | Antifungal | **C15** : $C_{71}H_{108}N_{12}O_{20}$<br>**C16** : $C_{72}H_{110}N_{12}O_{20}$<br>**C17** : $C_{73}H_{112}N_{12}O_{20}$<br>**C18** : $C_{74}H_{114}N_{12}O_{20}$<br>**C19** : $C_{75}H_{116}N_{12}O_{20}$ | 1449.7806 (4,8)<br>1463.8004 (1,9)<br>1477.8171 (2,5)<br>1491.8305 (2,7)<br>1505.8440 (4,1) | | |
| 4 Difficidins | Antibacterial | **Difficidin** : $C_{31}H_{45}O_6P$<br>**Oxydifficidin** : $C_{31}H_{45}O_7P$ | 567.2828 (3,2)<br>583.2795 (7,8) | | |
| 5 Bacillaenes | Antibacterial | **Bacillaene** : $C_{34}H_{48}N_2O_6$<br>**Dihydro-** : $C_{34}H_{50}N_2O_6$ | 581.3585 (2,3)<br>583.3726 (2,7) | | |
| 6 Macrolactins | Antibacterial | **A**:$C_{24}H_{34}O_5$(R=H)<br>***7-O-malonyl-A***:$C_{27}H_{36}O_8$ (R=CO-$CH_2$-COOH)<br>***7-O-succinyl-A***:$C_{28}H_{38}O_8$ (R=CO-$CH_2$-$CH_2$-COOH)<br>**D**:$C_{34}H_{48}O_{13}$ (R=6-*O*-syccinyl-β-gluc.) | 425.2298 (6,7)<br>511,2292 (2,0)<br>525,2459 (0,0)<br>687.2991 (0,6) | | |
| 7 Bacillibactin | Siderophore | $C_{39}H_{42}N_6O_{18}$ | 883.2620 (0,1) | | |
| 8 Orfamides | Antifungal ISR | **B** : $C_{63}H_{112}N_{10}O_{17}$<br>**D** : $C_{61}H_{108}N_{10}O_{17}$<br>**E** : $C_{63}H_{110}N_{10}O_{17}$ | 1281.8387 (8,4)<br>1253.8054 (7,0)<br>1279.8157 (2,6) | | |
| 9 Sessilins | Antifungal ISR | **A**:$C_{96}H_{166}N_{22}O_{25}$(R=Hse)<br>**B**:$C_{96}H_{168}N_{22}O_{26}$(acyclic A)<br>**C** :$C_{94}H_{162}N_{22}O_{24}$(R=Gly) | 1014.6457 (18,4)<br>1023.6473 (14,7)<br>992.6264 (12,6) | | |
| 10 Phenazines | Antibiotic ISR | **PCA** : $C_{13}H_8N_2O_2$ (R=OH)<br>**PCN** : $C_{13}H_9NO_3$ (R=$NH_2$) | 225.0681 (10,0)<br>224.0814 (2,0) | | |
| 11 Pyochelins | Siderophore | **Enantio-** : $C_{14}H_{16}N_2O_3S_2$ (4'S,2''S,4''S) | 325.0665 (3,1) | | |
| 12 Pyoverdine | Siderophore | $C_{55}H_{82}N_{15}O_{21}$ | 1289.5928 (3,5) | | |

**b**

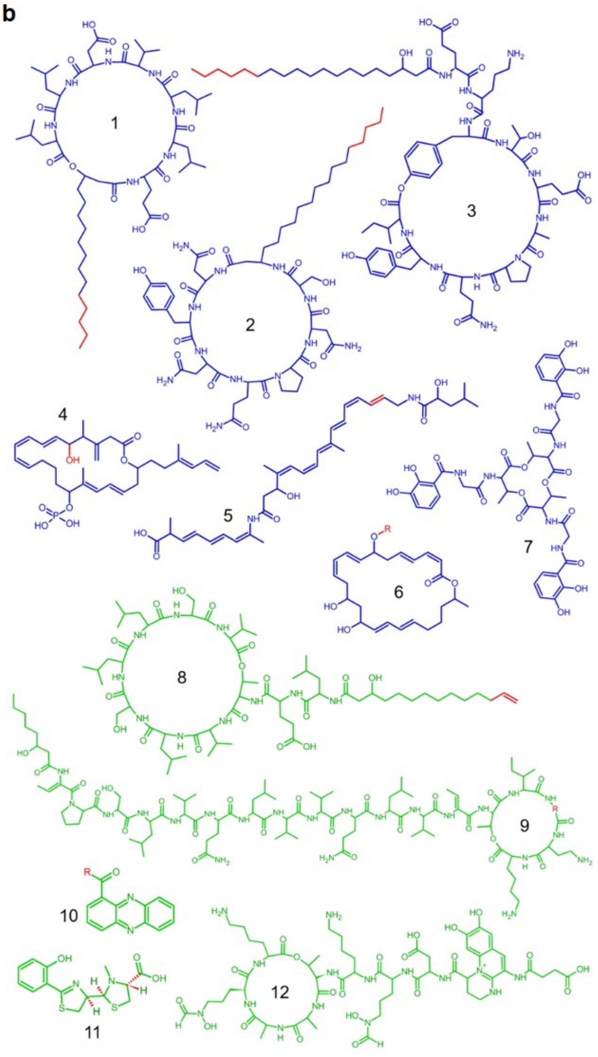

**Supplementary Figure 1. Chemical diversity in bioactive secondary metabolites produced by *B. velezensis* GA1 (blue) and *Pseudomonas* sp. CMR12a (green). a,** BSMs (numbers refer to the corresponding structure depicted in **b)**, their main described activities (ISR, induction of systemic resistance in plants) and the raw formula of detected variants with molecular mass and LC-MS chromatograms. BSMs were analyzed upon growth of the bacteria in exudate mimicking medium (EM) and casamino acid medium (CAA) **b,** BSMs structures with variable parts in red, explaining the natural co-production of variants for most of the compounds.

a

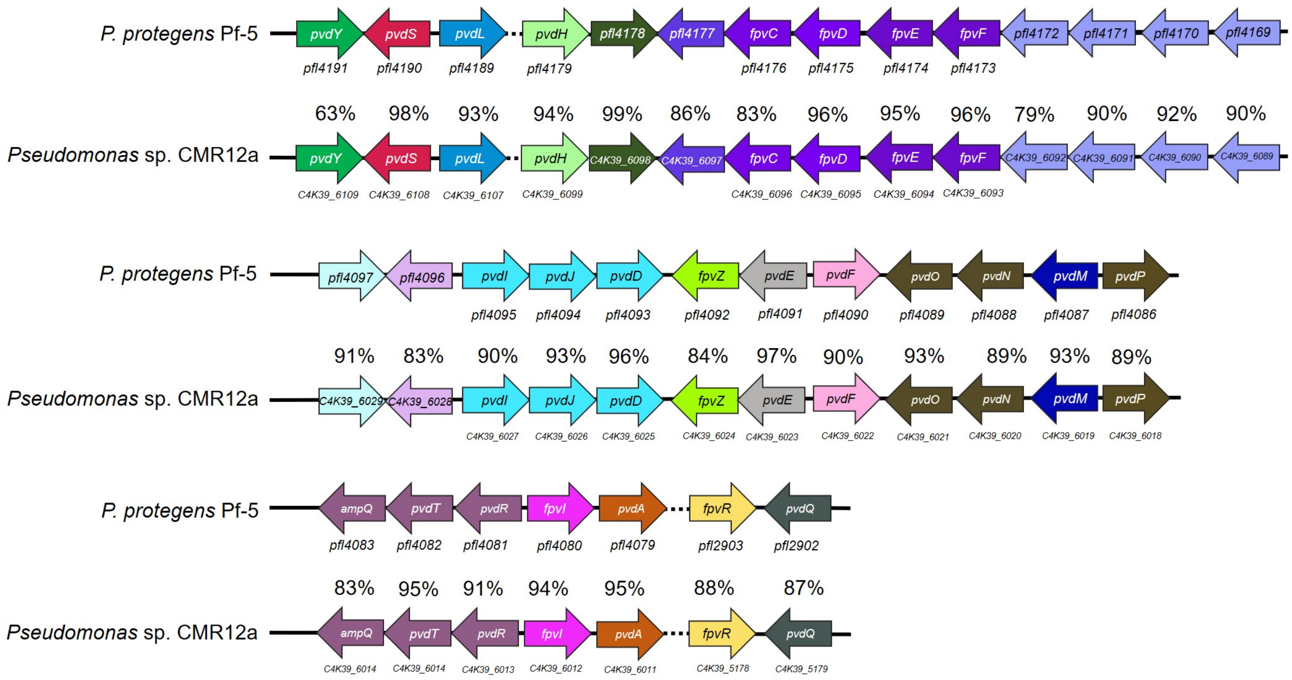


b

| FRAGMENTS | m/z | FRAGMENT INTENSITY |
|---|---|---|
| 1 | 1260.5857 | 2980.48 |
| 2/10 | 1270.5686 | 2025.38 |
| 3 | 759.2935 | 1620.38 |
| 4 | 648.2229 | 944.03 |
| 5 | 603.2024 | 829.93 |
| 6 | 490.1537 | 716.68 |
| 7 | 473.1280 | 21347.25 |
| 8 | 375.1266 | 2453.58 |
| 9 | 358.1015 | 1312.32 |
| 10 | 1270.5686 | 2025.38 |

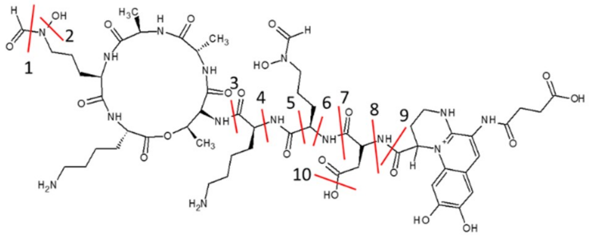


**Supplementary Figure 2**. ***Pseudomonas* sp. CMR12a produces a PVD structurally similar to the one of *P. protegens* Pf-5. a,** *Pseudomonas* sp. CMR12a genes were compared to the previously described *P. protegens* Pf-5 PVD biosynthetic cluster[1]. The corresponding locus tags are indicated for each gene. The nucleotide identity was calculated by blast in MAGE platform (https://mage.genoscope.cns.fr). **b,** The assigned fragments of *Pseudomonas* sp. CMR12a PVD confirms the structural similarity with the PVD from *P. protegens* Pf-5[1].

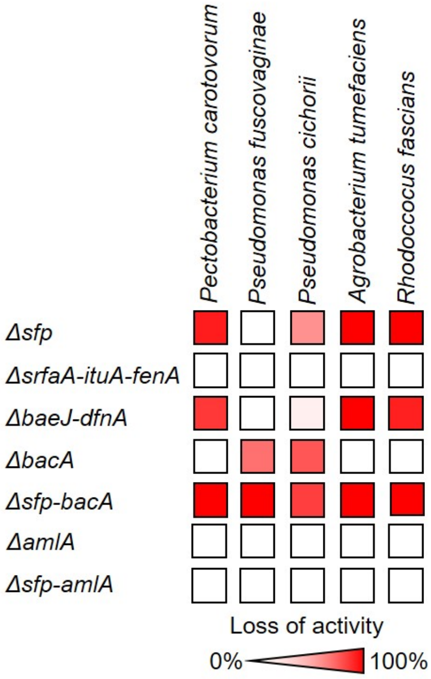


**Supplementary Figure 3. The anti-bacterial activity of *B. velezensis* GA1 relies on the production of different BSMs according to the target species.** The heatmap shows the loss of anti-bacterial activity of GA1 mutants compared to the WT strain when tested against five plant pathogenic species. The mutants are impaired in the production of NRPs and PKs (Δ*sfp*), surfactins, iturins and fengycins (Δ*srfaA-ituA-fenA*), difficidins and bacillaenes (Δ*dfnA-baeJ*), bacilysin (Δ*bacA*), NRPs, PKs and bacilysin (Δ*sfp-bacA*), amylolysin (Δ*amlA*) and NRPs, PKs and amylolysin (Δ*sfp-amlA*). The intensity of activity loss is represented by the color scale where the darkest red indicates loss of 100 % and white reflects no difference compared to the wild type. None of the mutants displayed gain in activity compared to the WT strain. Heatmap shows the mean of three biological replicates (n = 3).

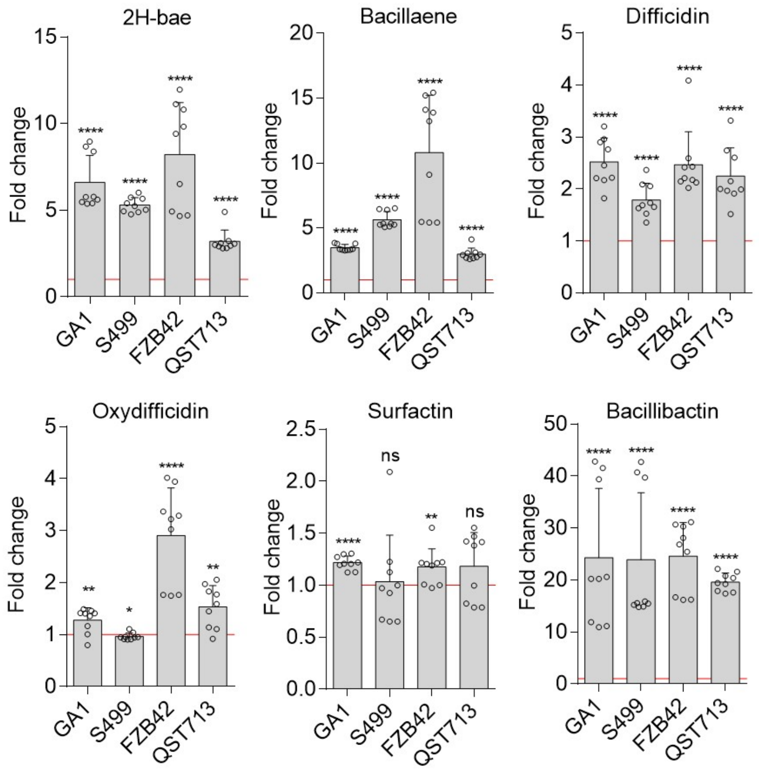


**Supplementary Figure 4. Effect of *Pseudomonas* sp. CMR12a metabolites on BSMs production by *B. velezensis* strains GA1, S499, FZB42 and QST713.** Data indicate fold increase in BSMs production upon addition of CMR12a CFS (4% v/v) compared to un-supplemented cultures (fold change = 1, red line). Data were calculated based on relative quantification of the compounds by UPLC-MS (peak area) in both conditions. Mean values and SD were calculated from three cultures (repeats) in three independent experiments (n = 9). Statistical significance was calculated using Mann–Whitney test where “ns” means no significant difference; “*”, $P<0.05$; “**”, $P<0.01$; “***”, $P<0.001$; “****”, $P<0.0001$.

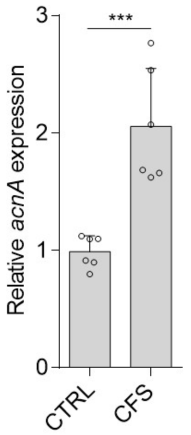


**Supplementary Figure 5. Stimulation of amylocyclicin in *B. velezensis* strain S499 in response to *Pseudomonas* metabolites.** Relative xpression of the *acnA* gene measured after 8 h of incubation in presence of *Pseudomonas* cell-free supernatant (CFS) (added at 2% v/v) compared to un-supplemented cultures (CTRL). Data show means and SD calculated from three cultures in two independent experiments (n = 6). The statistical difference, in *acnA* expression, between the two conditions was calculated by using T-test, "***", *P*<0.001.

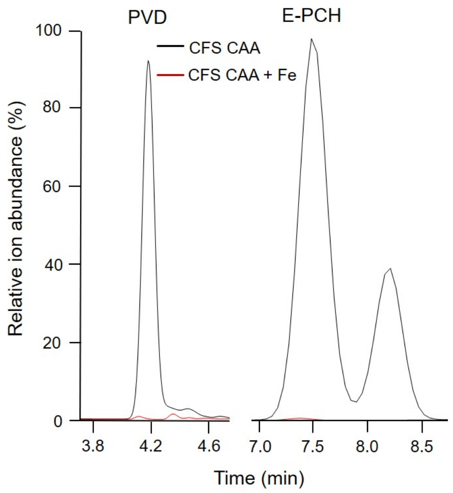


**Supplementary Figure 6. Addition of iron into the culture medium represses siderophore production by *Pseudomonas* sp. CMR12a.** UPLC-MS extracted ions chromatograms illustrating the relative abundance of ions corresponding to PVD (m/z 1289.5928) and E-PCH (m/z 325.0665) and as produced upon growth in CAA medium (black line) or in the medium supplemented with 20 µg/L of $FeCl_3.6H_2O$ (red line). The major peak at RT 3.95 min represents the main PVD form described in Supplementary Figure 2 while the minor peak at RT 4.2 min corresponds to a structural variant with similar peptide moiety but most probably a different side-chain in the chromophore (not identified).

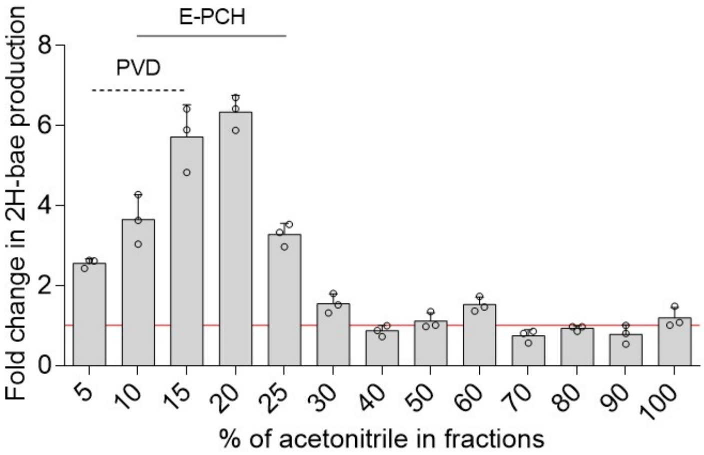


**Supplementary Figure 7. CFS fractions containing E-PCH are the most efficient at triggering 2H-bae production by *B. velezensis* GA1.** The CMR12a CFS was submitted to solid-phase extraction on C18 cartridges and compounds were stepwise eluted according to their hydrophobicity with increasing acetonitrile-water ratio, expressed in % (v/v). All fractions were analyzed by UPLC-MS and those containing substantial amounts of PVD and/or E-PCH are labeled as dashed and solid lines, respectively. The most active fractions 15% and 20% are the most enriched in E-PCH and fraction 10% contained the highest amounts of PVD. Fold change equal to 1 is represented as a red line and corresponds to the production of 2H-bae in *B. velezensis* GA1 un-supplemented culture. Data are from one representative experiment show mean and SD calculated from three technical replicates.

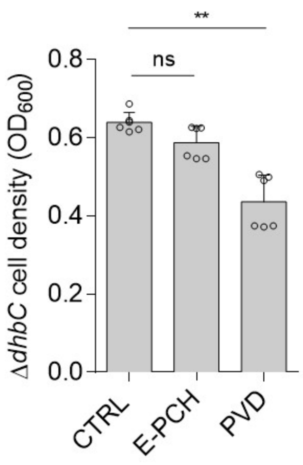


**Supplementary Figure 8. Effect of PVD and E-PCH on the growth of the bacillibactin-suppressed mutant of *B. velezensis* GA1.** Pure compounds were added to the *Bacillus* culture at a concentration corresponding to the one resulting from the addition of CFS CAA at 4% v/v. $OD_{600nm}$ was measured at the mid-exponential phase. Data are means and SE calculated from three replicate cultures in two independent experiments (n = 6), T-test with 'ns' = no significant difference; "**", statistically different at $P<0.01$.

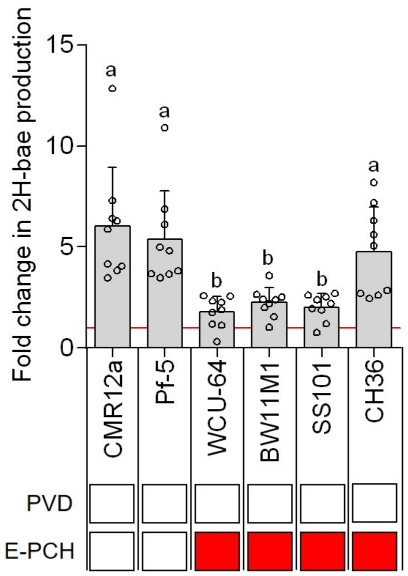


**Supplementary Figure 9: Stimulation of polyketide production by various *Pseudomonas* strains.** The 2H-bae production by *B. velezensis* GA1 is differently impacted by *P. protegens* Pf-5, WCU-84, *P. putida* BW11M1, *P. lactis* SS101 and *P. tolaasii* CH36, and is dependent on the production of PVD and E-PCH. *Bacillus* culture was supplemented with 4% (v/v) of *Pseudomonas* CFSs. Fold change equal to 1 is represented as a red line and corresponds to the production of 2H-bae in *B. velezensis* GA1 un-supplemented culture. Siderophore production or lack thereof is represented for each strain by white and red boxes respectively. The graph shows the mean and SD of three biological replicates with three technical repetitions (n = 9). Different letters indicate groups of statistically different conditions (one-way ANOVA and Tukey test; $P<0.05$). Production of metabolites in *B. velezensis* GA1 un-supplemented culture is not statistically different from the conditions within group b.

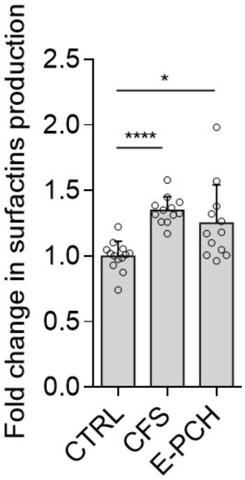


**Supplementary Figure 10. Stimulation of surfactins production in GA1 by E-PCH.** Fold increase in surfactins production measured upon supplementation of GA1 culture with either CMR12a CFS added at 4% (v/v) or pure E-PCH at a final concentration of 1.4 µM. The graph shows mean and SD calculated from four different assays involving three technical replicates for each treatment (n = 12). Statistical comparisons between treatments were realized with Mann–Whitney-test. “*”, $P<0.05$; “****”, $P<0.0001$.

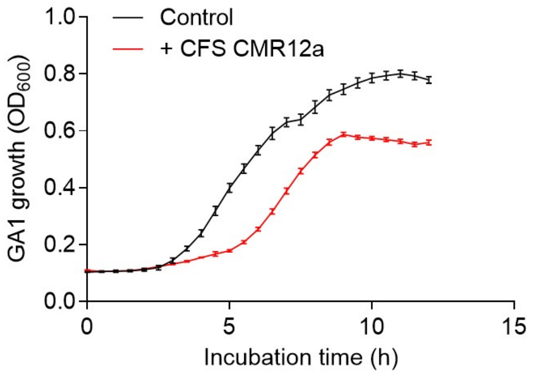


**Supplementary Figure 11. Effect of *Pseudomonas* sp. CMR12a metabolites on *B. velezensis* GA1 growth**: *B. velezensis* GA1 growth inhibition upon supplementation of culture medium with 4% v/v *Pseudomonas* CFS. Data show the mean and SD from three replicates.

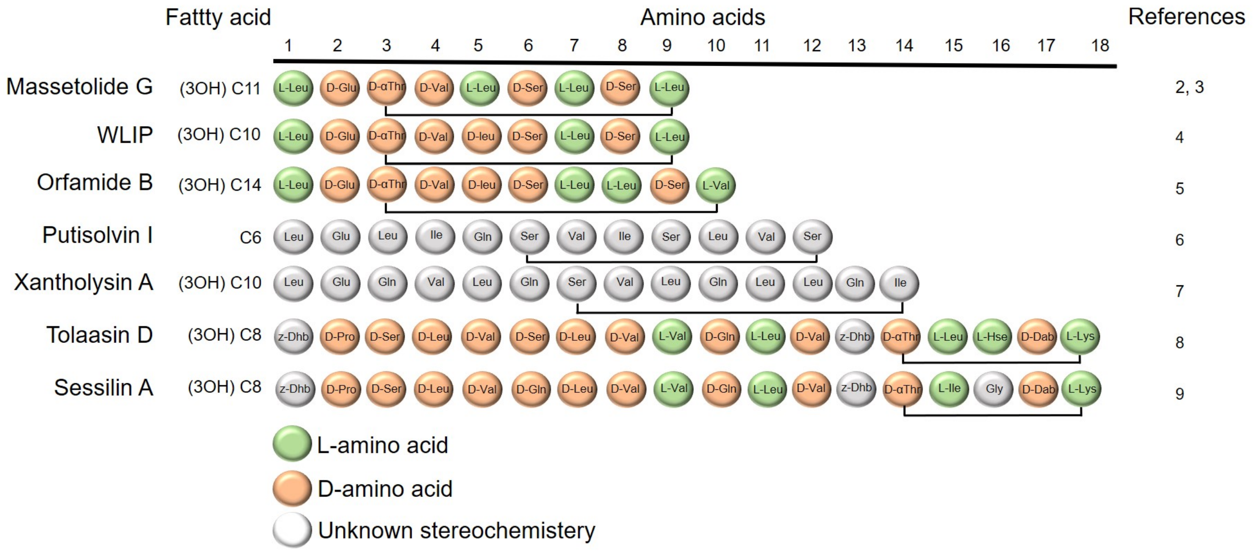


**Supplementary Fig. 12**. **Simplified structural representation of the cyclic lipopeptides produced by the different *Pseudomonas* strains used in this study**. The fatty acid chain length and the amino acid composition are depicted indicating where cyclization occurs. For each CLP, only the main form detected is represented, see Supplementary Figure 13 for UPLC-MS data and for more details about minor variants see also the corresponding associated references[2–7]).

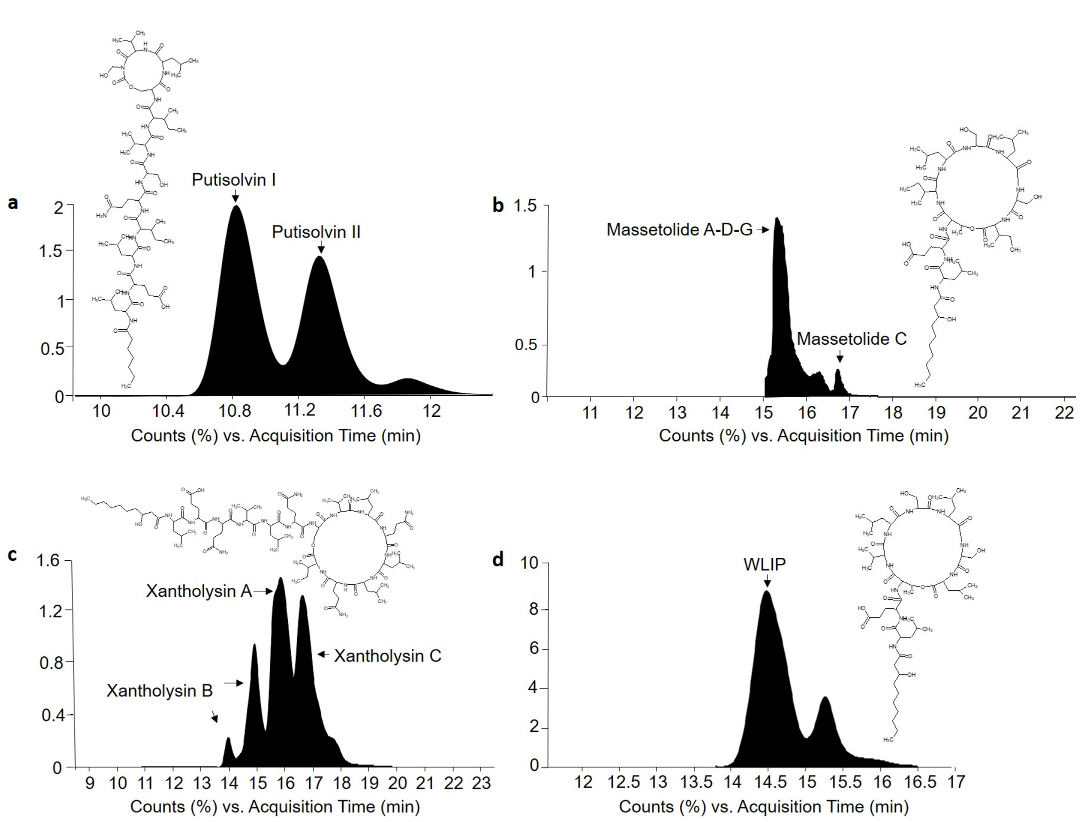


**Supplementary Figure 13. Main CLPs produced by *P. putida* WCU-84, *P. lactis* SS101, *P. putida* BW11M1 and *P. putida* RW10S2. a,** Relative ion abundance of putisolvin I and II produced by *P. putida* WCU-84. The structural formula of putisolvin I is presented. **b,** Relative ion abundance of massetolide A-D-G, C produced by *P. lactis* SS101. The structural formula of massetolide A-D-G is indicated. **c,** Relative ion abundance of xantholysin A, B and C produced by *P. putida* BW11M1. The structural formula of xantholysin A is presented. **d,** Relative ion abundance of WLIP produced by *P. putida* RW10S2. The structural formula of WLIP is presented.

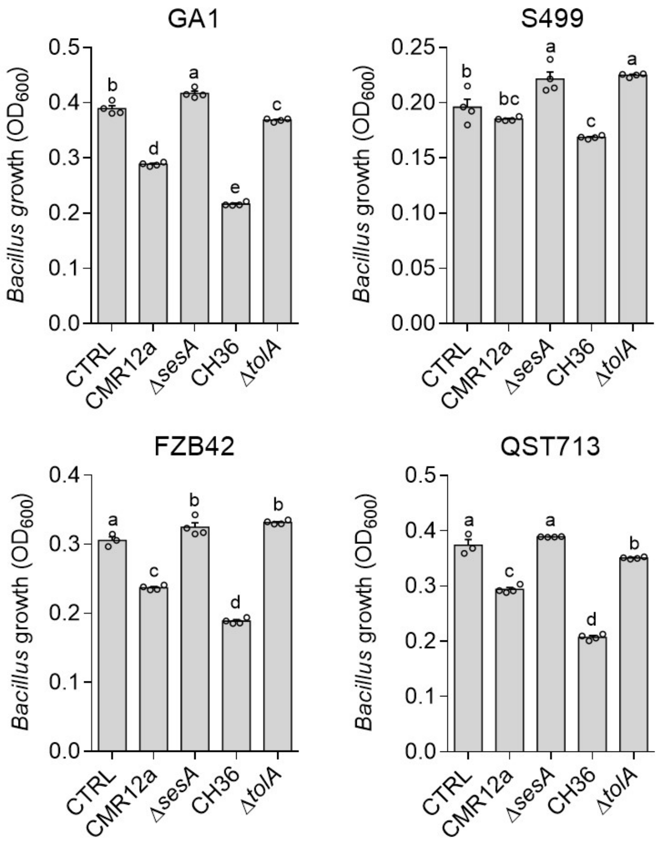


**Supplementary Figure 14. Sessilins and tolaasin inhibit the growth of different *B. velezensis* strains (FZB42, S499, QST713 and GA1).** Data show growth inhibitory effects of CFS from CMR12a WT or Δ*sesA* mutant impaired in sessilins production, and of CFS from *P. tolaasii* CH36 or its Δ*tolA* mutant repressed in tolaasin synthesis on *B. velezensis* strains growth. Graphs show mean and SD calculated from four replicate cultures in one representative experiment (n = 4). Letters a to d indicate statistically significant differences according to Tukey's test ($\alpha$ = 0.05).

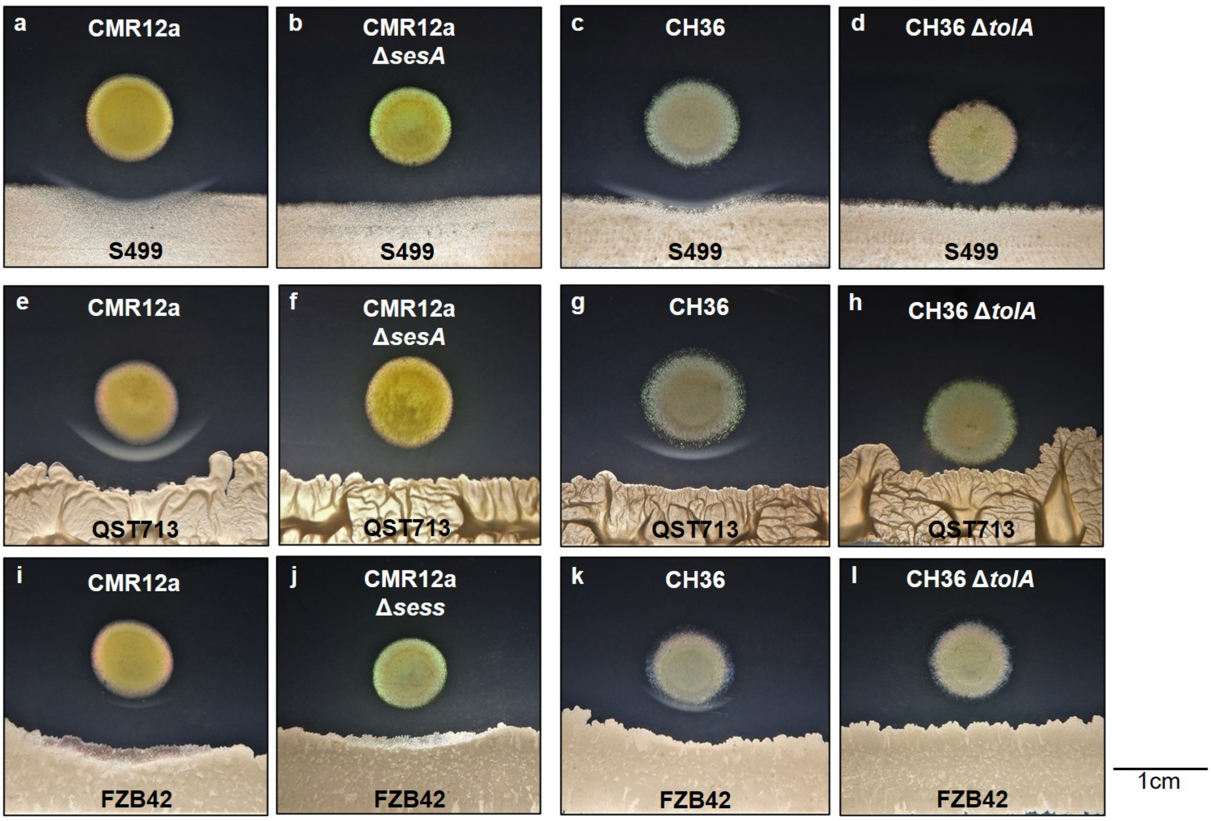


**Supplementary Figure 15. White-line formation between surfactins and sessilins/tolaasins is conserved within *B. velezensis* isolates.** The white line formation is dependent on the co-presence of surfactins and sessilins/tolaasins producers while during the interaction between *B. velezensis* WT strains and *Pseudomonas* sp. CMR12a (CMR12a) or *P. tolaasii* CH36 (CH36) mutants impaired in the production of sessilins or tolaasins, respectively, no white-line are observed. **a, c, e, g, i, k,** Sessilins/tolaasins production induces slight growth inhibition of all *B. velezensis.* **b, d, f, h, j, l,** No inhibition is observed in confrontation assays with *Pseudomonas* mutants unable to produce sessilins or tolaasins.

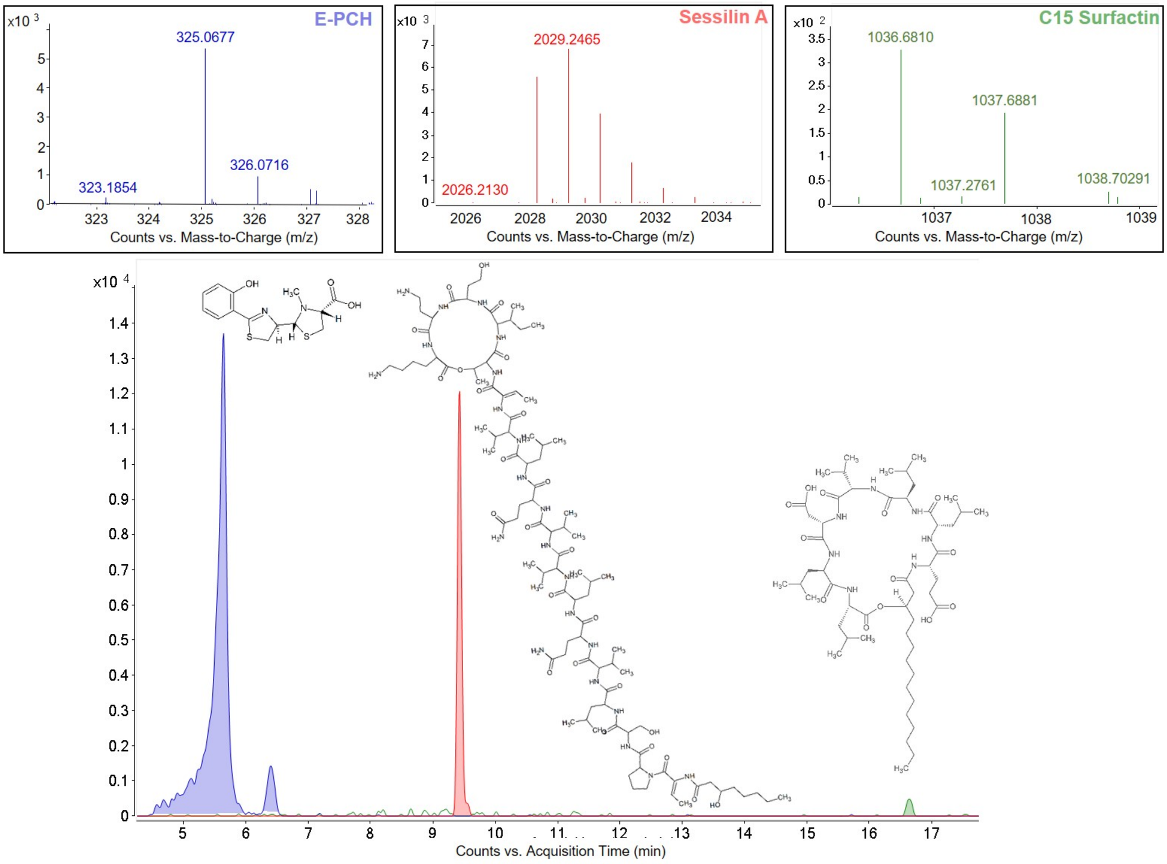


**Supplementary Figure 16. Quantification of sessilin, surfactin and E-PCH produced *in planta*.** LC-MS analysis of extracts from co-inoculated plants with *B. velezensis* GA1 and *Pseudomonas* sp. CMR12a after 3 days of co-inoculation on tomato plant roots. The chromatogram shows representative results from one of the two biological replicates obtained with at least four plants (n = 2).
